# GATK-gCNV: A Rare Copy Number Variant Discovery Algorithm and Its Application to Exome Sequencing in the UK Biobank

**DOI:** 10.1101/2022.08.25.504851

**Authors:** Mehrtash Babadi, Jack M. Fu, Samuel K. Lee, Andrey N. Smirnov, Laura D. Gauthier, Mark Walker, David I. Benjamin, Konrad J. Karczewski, Isaac Wong, Ryan L. Collins, Alba Sanchis-Juan, Harrison Brand, Eric Banks, Michael E. Talkowski

**Affiliations:** Data Sciences Platform, Broad Institute, Cambridge, MA, USA; Center for Genomic Medicine, Massachusetts General Hospital, Boston, MA, USA; Department of Neurology, Harvard Medical School, Boston, MA; Program in Medical and Population Genetics and Stanley Center for Psychiatric Research, Broad Institute, Cambridge, MA, USA; Analytic and Translational Genetics Unit, Massachusetts General Hospital, Boston, MA, USA; Stanley Center for Psychiatric Research, Broad Institute of MIT and Harvard, Cambridge, MA, USA

**Author notes:** These authors contributed equally (Mehrtash Babadi, Jack M. Fu, Samuel K. Lee, Andrey N. Smirnov).

**Keywords:** Copy Number Variation, Factor Analysis, UK Biobank, GATK, Terra, Autism, Structural Variation

## Abstract

Copy number variants (CNVs) are major contributors to genetic diversity and disease. To date, exome sequencing (ES) has been generated for millions of individuals in international biobanks, human disease studies, and clinical diagnostic screening. While standardized methods exist for detecting short variants (single nucleotide and insertion/deletion variants) using tools such as the Genome Analysis ToolKit (GATK), technical challenges have confounded similarly uniform large-scale CNV analyses from ES data. Given the profound impact of rare and *de novo* coding CNVs on genome organization and human disease, the lack of widely-adopted and robustly benchmarked rare CNV discovery tools has presented a barrier to routine exome-wide assessment of this critical class of variation. Here, we introduce GATK-gCNV, a flexible algorithm to discover rare CNVs from genome sequencing read-depth information, which we distribute as an open-source tool packaged in GATK. GATK-gCNV uses a probabilistic model and inference framework that accounts for technical biases while simultaneously predicting CNVs, which enables self-consistency between technical read-depth normalization and variant calling. We benchmarked GATK-gCNV in 7,962 exomes from individuals in quartet families with matched genome sequencing and microarray data. These analyses demonstrated 97% recall of rare (≤1% site frequency) coding CNVs detected by microarrays and 95% recall of rare coding CNVs discovered by genome sequencing at a resolution of more than two exons. We applied GATK-gCNV to generate a reference catalog of rare coding CNVs in 197,306 individuals with ES from the UK Biobank. We observed strong correlations between CNV rates per gene and measures of mutational constraint, as well as rare CNV associations with multiple traits. In summary, GATK-gCNV is a tunable approach for sensitive and specific CNV discovery in ES, which can easily be applied across trait association and clinical screening.

## INTRODUCTION

Copy number variants (CNVs) comprise duplications and deletions of genomic segments spanning ≥50 nucleotides. These gains and losses of genetic material can impact gene function and regulation with profound consequences in human disease^1,2^. While each human genome likely harbors more than 25,000 structural variants^3^, most gene-disrupting CNVs, including the vast majority of clinically interpretable pathogenic CNVs, experience strong negative selection and are therefore rare in the general population^4^. Thus, the ability to discover rare and *de novo* CNVs that alter protein-coding sequences with high recall and precision can have widespread utility in human genetic research, trait association, and clinical diagnostics.

The discovery of CNVs in many biomedical settings has historically relied upon low-resolution technologies like chromosomal microarray (CMA). Despite its technical limitations, exploration of large CNVs from CMA has provided substantial insights for many diseases. For example, CNV analysis using CMA is the first-tier diagnostic test recommended by the American College of Medical Genetics for children with unexplained developmental disorders^5^. However, the low resolution of genome-wide CMA precludes most gene- and exon-level interpretation of CNVs. Exome sequencing (ES) has revolutionized human disease research and diagnostic screening by enabling discovery of variation in protein-coding sequences while being substantially cheaper than genome sequencing (GS)^6,7^. In theory, ES should permit the detection of most CNVs that alter genes with equivalent recall to GS and represent a marked improvement in resolution beyond CMA in research and clinical settings. In practice, variability in sequencing coverage due to hybridization-based exome enrichment^8^ and other biases related to ES library preparation^9^ distort informative read-depth signals depending on both local sequence context and the properties of each individual sample. This technical variability has presented significant challenges in balancing recall of ES-based CNV discovery with the need for high precision in many applications, such as rare variant disease association studies or the accurate identification of *de novo* CNVs in clinical diagnostics. There are existing methods for CNV detection in ES that attempt to remove systematic biases and normalize read-depth data via PCA denoising^10^, regression^11^, pre-clustering of samples^12,13^, or accounting for genomic context such as GC-content^14^. CNVs are then detected in a second step using hidden Markov models (HMM) or nonparametric change-point detection algorithms applied to the normalized data^15^. These methods introduce a lack of self-consistency between the removal of systematic biases and the CNV calling by performing them in two separate steps, which can inadvertently remove informative CNV signals in the former and cause decreased recall in the latter.

The generation of ES data on millions of individuals to date^16–20^ provides a unique opportunity for large-scale assessment of rare CNVs across human diseases and traits. Whereas the use of the Genome Analysis Toolkit (GATK) to capture SNVs and indels in ES is well-established and ubiquitous^21^, the absence of a CNV discovery tool that can be routinely applied to ES data with comparable performance, documentation, and dedicated support to GATK’s functionality for SNV/indel analysis represents a significant barrier to realizing the full potential of ES data. Here, we present GATK-gCNV, a principled Bayesian method for learning global and sample-specific biases of read-depth data from large cohorts while simultaneously detecting CNVs. Our model combines a negative-binomial factor-analysis module for learning genome-wide latent factors of technical read-depth variation together with a hierarchical hidden Markov model (HHMM) for detecting singleton and rare polymorphic CNVs in ES cohorts. In addition to being packaged as part of the GATK, we also provide GATK-gCNV as a cloud-enabled tool in the Terra cloud platform (https://terra.bio) for easy adoption by the biomedical community. We provide extensive benchmarking of GATK-gCNV against gold-standard GS and CMA data in autism quartet families harboring pathogenic CNVs, and we demonstrate the scalable utility of GATK-gCNV by generating a reference map of rare genic CNVs in 197,306 ES samples from the UK Biobank. From these data, negative selection against coding loss-of-function (LoF) variants was strongly correlated with the rates of rare deletions and duplications of individual genes. We also examined rare CNV trait associations in the UKBB. These results highlight that rare gene-disruptive variation that is accessible to ES data can be routinely captured at very large-scale for low cost in ES-based association studies and diagnostic screening.

## RESULTS

### Algorithm overview

We developed an algorithm, GATK-gCNV, to jointly discover and genotype CNVs across ES datasets using read-depth information (**Fig. 1**). While GATK-gCNV can and has been optimized for similar analyses in GS datasets, the analyses presented here focus on ES methods and applications where technical sources of read-depth variation pose a major hurdle to CNV detection. The algorithm is summarized here and provided in complete detail in **Online Methods** and **Supplementary Note**.

**Fig. 1.**
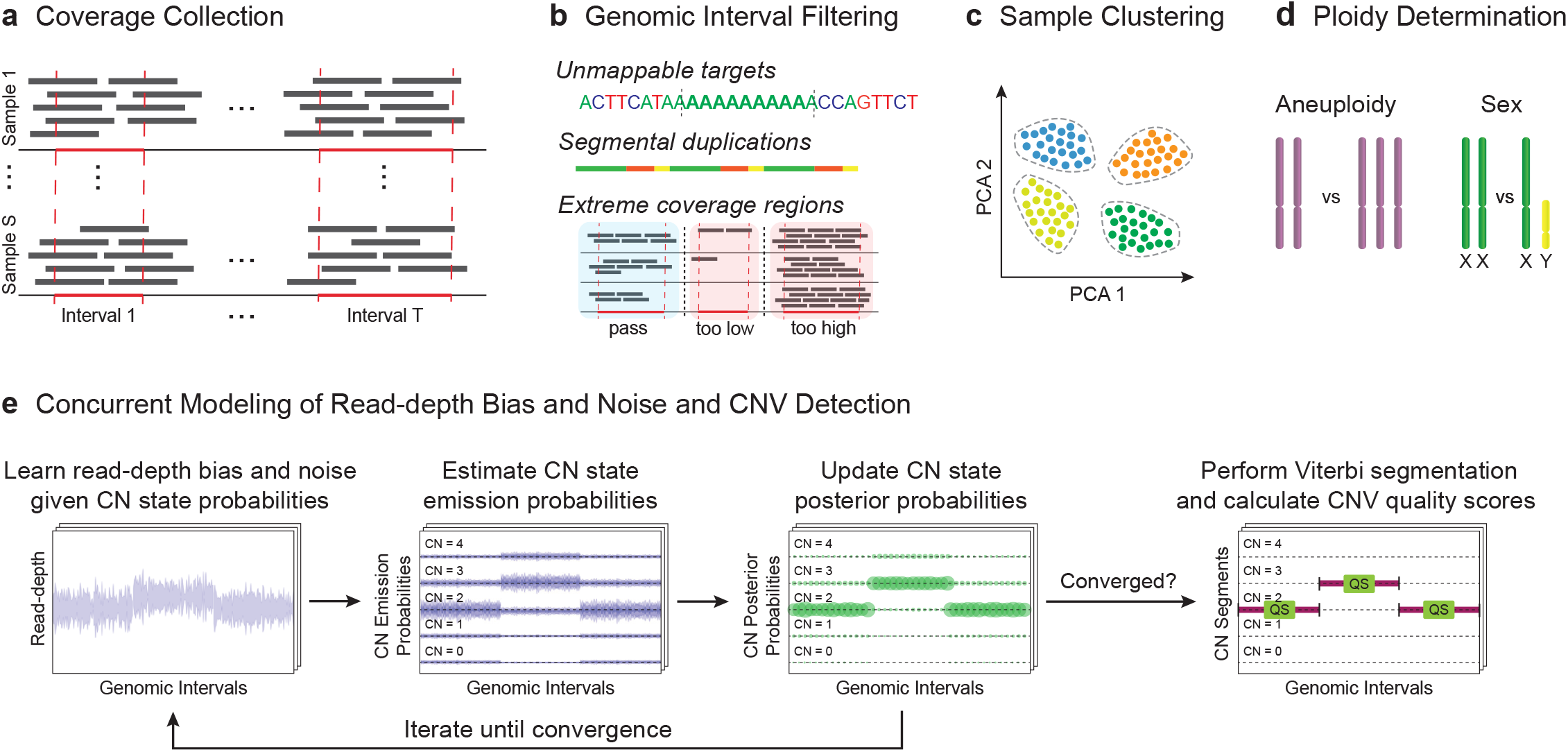
GATK-gCNV pipeline steps. **a**, Coverage information is collected from genome-aligned reads over a set of predefined genomic intervals. **b**, The original interval list is filtered to remove coverage outliers, unmappable genomic sequence, and regions of segmental duplications. **c**, Samples are clustered into batches based on read-depth profile similarity and each batch is processed separately. **d**, Chromosomal ploidies are inferred using total read-depth of each chromosome. **e**, The GATK-gCNV model learns read-depth bias and noise and iteratively updates copy number state posterior probabilities until a self-consistent state is obtained; after convergence, constant copy number segments are found using the Viterbi algorithm along with segmentation quality scores. **Abbreviations**: CN - copy number; QS - quality score.

GATK-gCNV begins by calculating read counts over user-defined target regions (e.g., exons) in each sample while excluding regions with problematic sequence content. Next, samples with technically similar read-depth profiles are clustered into batches via principal components analysis (PCA) to reduce technical biases and improve computational efficiency during processing. After clustering, the ploidy (i.e., copy number) of every chromosome is estimated for each sample to detect potential aneuploidies and determine sample sex. After all data has been preprocessed, GATK-gCNV performs read-depth denoising and CNV inference within a unified probabilistic model and determines CNV boundaries via the Viterbi algorithm (**Supplementary Fig. 1**). In practice, GATK-gCNV can be executed in two modes: “cohort” mode and “case” mode. Cohort-mode uses all input samples to train a read-count model while simultaneously inferring CNVs in each sample, whereas case-mode applies a pre-trained model to call CNVs for any number of additional samples in parallel. Generating CNV calls through case-mode is much faster and computationally cheaper, as it avoids the costly step of training a new read-count model. To take full advantage of both modes, we subsample up to 200 of each PCA-defined batch to run cohort-mode, then apply case-mode to the remaining samples in each batch, greatly saving on cost.

### Benchmarking GATK-gCNV performance using 7,962 deeply-profiled genomes

We assessed GATK-gCNV performance to discover rare and *de novo* CNVs on ES data (∼75x coverage)^22^ from the Simons Simplex Collection (SSC), which is a cohort of autism spectrum disorder (ASD) families that have previously undergone CNV detection and rigorous quality control with CMA (2,591 families^23^) and GS (2,672 families^24^) data. In total, we assessed 7,962 ES samples with matched GS data^24–26^ and 7,636 samples with matched CMA data^23^, both of which provided ground-truth. The family-based structure of the SSC also enabled the assessment of both rare and *de novo* CNVs at the resolution of individual genes and exons across 3,131 parent-child trios (1,208 families contributed multiple trios). To measure the performance of GATK-gCNV, we calculated the recall and precision compared to GS-or CMA-based CNV datasets. When assessing recall, we defined a CNV from GS or CMA to be captured by GATK-gCNV if at least 50% of well-captured intervals (defined below) spanning the variant were overlapped by ES CNV predictions in at least 50% of the same samples. For precision, we deemed a GATK-gCNV variant to have GS support if 50% of the well-captured intervals of that variant were overlapped by a matching GS CNV called in at least 50% of the same samples.

We applied GATK-gCNV to all SSC samples using the cloud-based Terra platform for biomedical research (http://terra.bio/) and have deployed a demonstration workspace of GATK-gCNV as a resource for the community (**Online Methods**). We implemented PCA-based sample batching based on a set of 7,981 curated intervals that differentiated common ES capture technologies, sequencing centers, and other technical factors (**Fig. 2a,b**, **Online Methods**). This approach subdivided the SSC ES samples into 14 batches of approximately 722 samples each (Interquartile range [IQR]=466; **Fig. 2c**). To further harmonize different exon-capture targets and bait sets across studies, we restricted all analyses to protein-coding exons from canonical transcripts as described in GENCODE v33^27^ and merged overlapping regions to derive a consensus set of 190,488 autosomal exons. We additionally filtered out regions of extreme GC-content, repetitive sequence content, and poor mappability, and subdivided large exons to produce a final set of 330,526 intervals for CNV discovery (median size=384 bp, IQR=518; **Online Methods**). Within each PCA-defined batch of samples, we further filtered intervals based on low sequencing coverage (median <10 reads per sample). On average, this batch-specific coverage filtering resulted in 187,804 (IQR=55,732) intervals retained for CNV discovery per batch, corresponding to 169,442 exons on average (IQR=17,492). Hereafter, we refer to these intervals as “well-captured”.

**Fig. 2.**
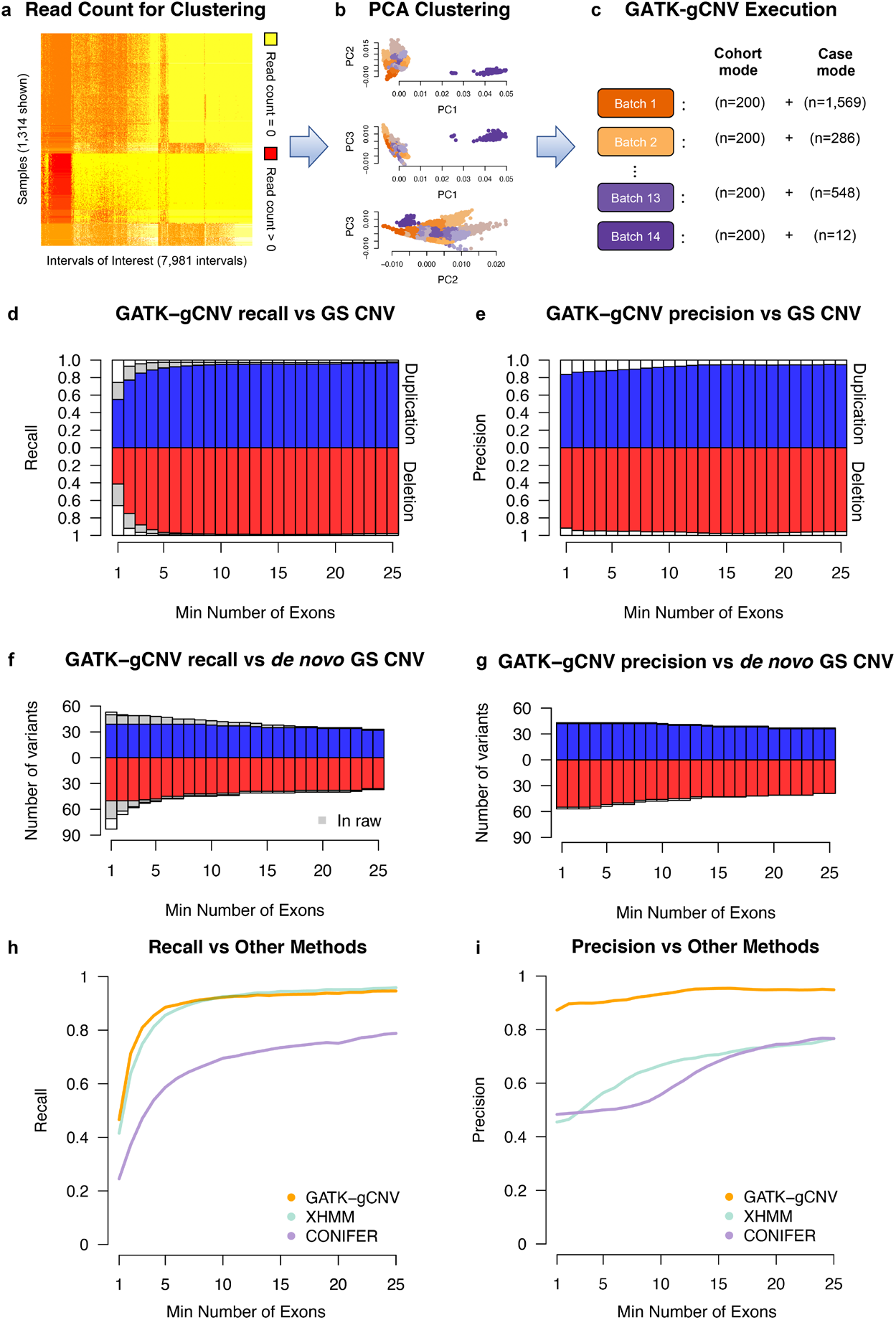
Calling and benchmarking of GATK-gCNV callset in a cohort of more than 7,000 samples with matching deep GS sequencing. **a**, A heatmap illustration of the distinct read-count signal of the 7,981 intervals chosen for the batch creation procedure. **b**, After normalizing for median read count, the first three PCs are clustered to determine which samples will be processed together with GATK-gCNV, colored by the assigned batch. **c**, For each of the 14 batches generated, a random subset of 200 samples was chosen to generate a read-count model using cohort-mode; the remaining samples were processed in case-mode. **d**, The recall (and **e**, precision) of rare CNVs in GATK-gCNV ES CNVs compared to GS gold-standard CNVs as a function of the number of exons the variant spans. **f**, The recall (and **g**, precision) of de novo CNVs in GATK-gCNV compared to gold-standard GS CNVs as a function of the number of exons. **h**, The recall (and **i**, precision) of rare CNVs in GATK-gCNV, XHMM, and CONIFER ES CNVs compared to GS gold-standard CNVs as a function of the number of exons the variant spans. **Abbreviations**: PCA - principal component analysis; ES-exome sequencing; GS - whole genome sequencing.

We executed GATK-gCNV in cohort-mode on random subsets of 200 samples from each PCA-defined batch, training a CNV-discovery model tailored to the specific technical properties of each batch. GATK-gCNV cohort-mode required a median of 9:05 hours wall clock time to train and call each batch, processing 12,500 intervals at a time in parallel compute instances with 4 CPU cores and 24GB memory total. When run in this configuration on Terra, cohort-mode cost $0.037 per sample. After training a CNV discovery model for each batch, we conducted CNV discovery on all remaining samples per batch using GATK-gCNV case-mode, which required a median of 7:42 hours wall clock time and $0.021 per sample on Terra. By leveraging the highly parallelized computing possible on cloud-based platforms like Terra, we were able to process 7,962 SSC samples in less than 24 hours of wall time and at $0.026 per sample.

By design, the unfiltered output of GATK-gCNV is extremely sensitive to allow for exhaustive searches of candidate CNVs, producing an average of 6.3 rare (variant site frequency < 1%) CNV calls per sample (2.4 deletions and 3.9 duplications) at a resolution of more than two well-captured exons. At this resolution, the raw GATK-gCNV output achieved 95% recall in 7,962 SSC samples with matching ES and GS data (**Supplementary Fig. 2a-d, Table 1**), but precision is low (22%) when using unfiltered outputs. We developed a series of sample- and variant-level filters to define high-confidence CNVs for applications where high precision is critical, such as trait association studies or *de novo* CNV prediction. For variant-level filtering, we leveraged a quality metric (QS) emitted by GATK-gCNV for each CNV, which models the probability in Phred scale that at least one interval within the CNV event locus was consistent with the estimated copy number state. We assigned a dynamic minimum QS threshold that scales with increasing CNV size, and fit this threshold separately for each combination of CNV type (deletion or duplication) and zygosity (heterozygous or homozygous), as described in **Online Methods**. For sample-level filtering, we found that the total number of CNV calls per sample correlated with the overall reliability and calibration of that sample, and that thresholds of >200 raw CNV calls or >35 CNVs with QS>20 were able to isolate poor-quality samples.

**Table 1.**
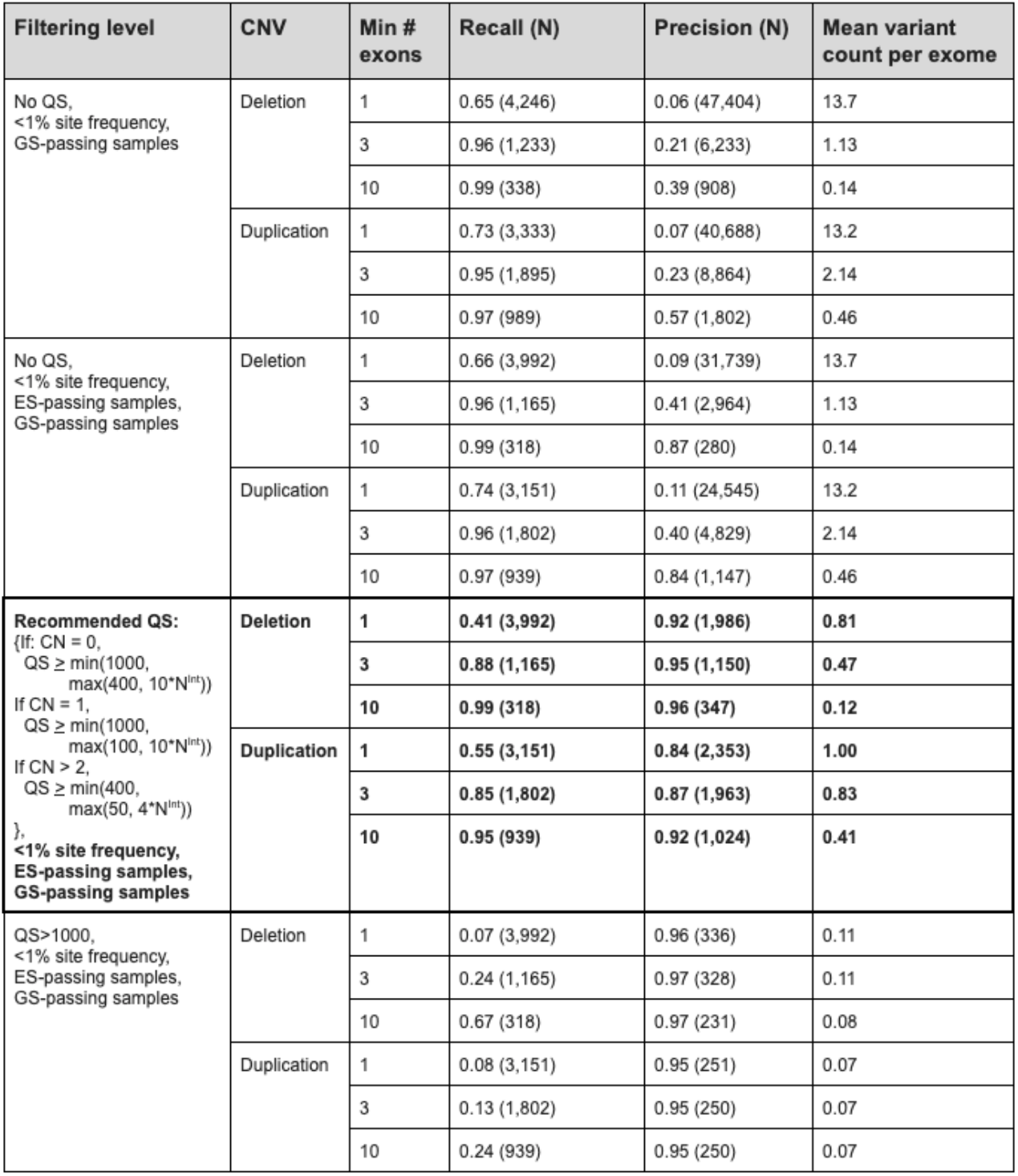
Performance of GATK-gCNV at different filtering thresholds. Various filtering thresholds can be adopted for the desired GATK-gCNV performance. **Abbreviations**: QS - quality score output by GATK-gCNV used for call-level filtering; GS - genome sequencing; ES - exome sequencing; CN - copy number; N^Int^ - number of well-captured intervals.

| Filtering level | CNV | Min # exons | Recall (N) | Precision (N) | Mean variant count per exome |
| --- | --- | --- | --- | --- | --- |
| No QS,<br><1% site frequency,<br>GS-passing samples | Deletion | 1 | 0.65 (4,246) | 0.06 (47,404) | 13.7 |
|  |  | 3 | 0.96 (1,233) | 0.21 (6,233) | 1.13 |
|  |  | 10 | 0.99 (338) | 0.39 (908) | 0.14 |
|  | Duplication | 1 | 0.73 (3,333) | 0.07 (40,688) | 13.2 |
|  |  | 3 | 0.95 (1,895) | 0.23 (8,864) | 2.14 |
|  |  | 10 | 0.97 (989) | 0.57 (1,802) | 0.46 |
| No QS,<br><1% site frequency,<br>ES-passing samples,<br>GS-passing samples | Deletion | 1 | 0.66 (3,992) | 0.09 (31,739) | 13.7 |
|  |  | 3 | 0.96 (1,165) | 0.41 (2,964) | 1.13 |
|  |  | 10 | 0.99 (318) | 0.87 (280) | 0.14 |
|  | Duplication | 1 | 0.74 (3,151) | 0.11 (24,545) | 13.2 |
|  |  | 3 | 0.96 (1,802) | 0.40 (4,829) | 2.14 |
|  |  | 10 | 0.97 (939) | 0.84 (1,147) | 0.46 |
| **Recommended QS:**<br>{If: CN = 0,<br>QS ≥ min(1000,<br>max(400, 10*N<sup>int</sup>))<br>If CN = 1,<br>QS ≥ min(1000,<br>max(100, 10*N<sup>int</sup>))<br>If CN > 2,<br>QS ≥ min(400,<br>max(50, 4*N<sup>int</sup>))<br>},<br><1% site frequency,<br>ES-passing samples,<br>GS-passing samples | Deletion | 1 | 0.41 (3,992) | 0.92 (1,966) | 0.81 |
|  |  | 3 | 0.88 (1,165) | 0.95 (1,150) | 0.47 |
|  |  | 10 | 0.99 (318) | 0.96 (347) | 0.12 |
|  | Duplication | 1 | 0.55 (3,151) | 0.84 (2,353) | 1.00 |
|  |  | 3 | 0.85 (1,802) | 0.87 (1,963) | 0.83 |
|  |  | 10 | 0.95 (939) | 0.92 (1,024) | 0.41 |
| QS>1000,<br><1% site frequency,<br>ES-passing samples,<br>GS-passing samples | Deletion | 1 | 0.07 (3,992) | 0.96 (336) | 0.11 |
|  |  | 3 | 0.24 (1,165) | 0.97 (328) | 0.11 |
|  |  | 10 | 0.67 (318) | 0.97 (231) | 0.08 |
|  | Duplication | 1 | 0.08 (3,151) | 0.95 (251) | 0.07 |
|  |  | 3 | 0.13 (1,802) | 0.95 (250) | 0.07 |
|  |  | 10 | 0.24 (939) | 0.95 (250) | 0.07 |

Applying these *post hoc* filters in the SSC ES data retained 89% (7,116/7,962) of all SSC ES samples and generated a high-quality callset of 9,246 autosomal CNV calls corresponding to 3,119 unique variants spanning more than two well-captured exons, or an average of 1.3 CNVs per sample (0.47 deletions and 0.83 duplications). In this high-quality callset, deletions had a median size of 6 exons and duplications a median size of 10 exons, while 72% of samples carried at least one such CNV (37% carried a deletion, and 55% carried a duplication). Benchmarking these high-quality CNV calls against matched GS data revealed high precision (90%) with good recall (96% unfiltered, 86% after filtering) (**Fig. 2d,e, Table 1, Supplementary Fig. 3**). The QS threshold can be further raised for increased precision, where for example a threshold of QS>1000 produces precision of 96% for all variants, at the cost of reduced sensitivity (**Supplementary Fig. 2e,f, Table 1**). We also evaluated the performance of our high-quality GATK-gCNV callset versus rare CNVs (<1% site frequency) identified by CMA in 7,157 SSC samples for which we had matching ES and CMA data. After restricting to large (>50 kilobases & >2 exons), high-confidence CNVs from CMA (probability pCNV < 10^−9^ from Sanders *et al*.^23^), the high-quality GATK-gCNV callset achieved 97% recall of these CMA events (**Supplementary Fig. 4**). These benchmarks indicate that GATK-gCNV is likely to be sufficiently sensitive to displace CMA in diagnostic screening for protein-coding CNVs, with ES providing the added benefit of simultaneously capturing all coding SNVs and indels in that same sample.

We next benchmarked the accuracy of our GATK-gCNV pipeline in identifying *de novo* CNVs in the children of SSC families. We first predicted the transmission for each high-quality CNV identified in ES samples whose parents were both also present in our GATK-gCNV callset. This procedure identified 99 high-quality *de novo* CNVs (56 deletions and 43 duplications) among 3,097 children (mean = 0.032 *de novo CNVs* per child), which ranged in size from 3 to 667 exons (mean = 143 exons; median = 112 exons). We assessed the accuracy of these 99 *de novo* CNVs by directly comparing them to matched GS-derived *de novo* CNVs from the same samples^24^, finding that GATK-gCNV achieved 97% precision across all sizes of *de novo* CNVs, while maintaining 86% and 80% recall for 56 *de novo* deletions and 64 *de novo* duplications reported in the gold-standard GS dataset that spanned more than 2 well-captured exons respectively (**Fig. 2f,g**).

Lastly, we compared GATK-gCNV results to two existing CNV tools. XHMM leverages a PCA denoising step followed by an HMM based calling step and was used to generate the largest publicly available exome-derived CNV reference to date^4,28^. The other is CONIFER, which uses Singular Value Decomposition to normalize ES read-count variability followed by a threshold heuristic for CNV calling^29^. All evaluated CNV tools received as input the set of 330,526 intervals described above. We were able to process 96.3% (7,665/7,962 with accessible CRAMs) of the SSC samples with ES data using both XHMM and CONIFER in the same batches as in our GATK-gCNV implementation (**Online Methods**). Sample- and call-level filtering were conducted according to published best-practices for each method, including the removal of low-quality samples, intervals, and CNV calls. On average, out of the 190,488 non-overlapping exons, GATK-gCNV retained 169,442 exons (89%), XHMM retained 157,507 exons (83%), and CONIFER retained 136,957 exons (72%). Using the set of samples passing QC from each method and evaluating on the basis of all unfiltered, non-overlapping GENCODE v33 exons, GATK-gCNV achieved recall and precision of 81% and 90%, respectively; XHMM 75% and 50%, and CONIFER 47% and 49%, all at a resolution of >2 exons (**Fig. 2h,i**).

### A rare CNV resource of 197,306 UK Biobank participants

Following extensive benchmarking of GATK-gCNV, we demonstrated the utility of this method to uniformly process and obtain CNV calls in a recent disease association study of ASD^18^. Here, we demonstrate the scalability of the algorithm in application to biobank-scale datasets, where computational efficiency, cost, and performance are all important factors when conducting variant discovery.

The UK Biobank (UKBB)^30^ is one of the world’s largest population-based biobanks with ES data linked to deep electronic health information, including nearly 500,000 residents of the UK. At the time of these analyses, a total of 200,624 ES samples from the UKBB were available to the research community, which represented the largest collection of publicly accessible ES data. Several trait association studies have already been conducted from CMA and ES in these samples^16,31^. However, the patterns of rare coding CNVs in the UKBB at the resolution of individual exons and genes remain unknown. We therefore sought to demonstrate the utility of GATK-gCNV by generating a uniform, high-quality rare CNV resource from the UKBB to benchmark expectations for GATK-gCNV in similar large cohorts and as a novel CNV dataset for use by the biomedical research community.

We processed 200,624 UKBB exomes using GATK-gCNV with the method described above. Samples were clustered into 110 independent batches using the PCA-based clustering procedure described previously (median 1,687 samples per batch, IQR=1,300). We randomly selected 200 samples from each cluster to train a model with GATK-gCNV in the cohort-mode, with the remainder of samples in each cluster processed in case-mode using the cluster-matched model. We then applied the same sample- and variant-level filtering as used in the SSC cohort. The entire UKBB callset was processed in 60.05 hours of wall clock time. This was spread across 16,069 parallel CPU hours for 110 cohort-mode runs and 110 matching case-mode runs. The total cost to process all 200,624 samples was $6,423.44 ($0.032 per sample), including $1,002.43 for 22,000 samples in cohort-mode ($0.046 per sample) and $4,184.07 for 178,624 samples in case-mode ($0.023 per sample).

After applying all sample- and variant-level quality filters as described above, only 1.7% (3,318) of samples failed to meet our stringent sample-level thresholds. Across all 197,306 high-quality samples, we discovered 207,017 high-confidence rare CNV calls corresponding to 38,731 unique variants spanning >2 exons (**Fig. 3a**). Most samples (64%) carried at least one rare coding CNV that passed our filtering: 31% and 49% of samples carried at least one rare deletion and duplication of >2 exons, respectively (**Fig. 3b**). As expected, we found that coding deletions were smaller on average (median size: 6 exons) than coding duplications (median size: 12 exons), which likely reflects a combination of stronger purifying selection on large coding deletions^26^ and the comparatively higher technical difficulty for sensitive discovery of small duplications. We have returned these high-quality CNVs to the UKBB for dissemination to qualified researchers through the UKBB’s data-release procedure.

**Fig. 3.**
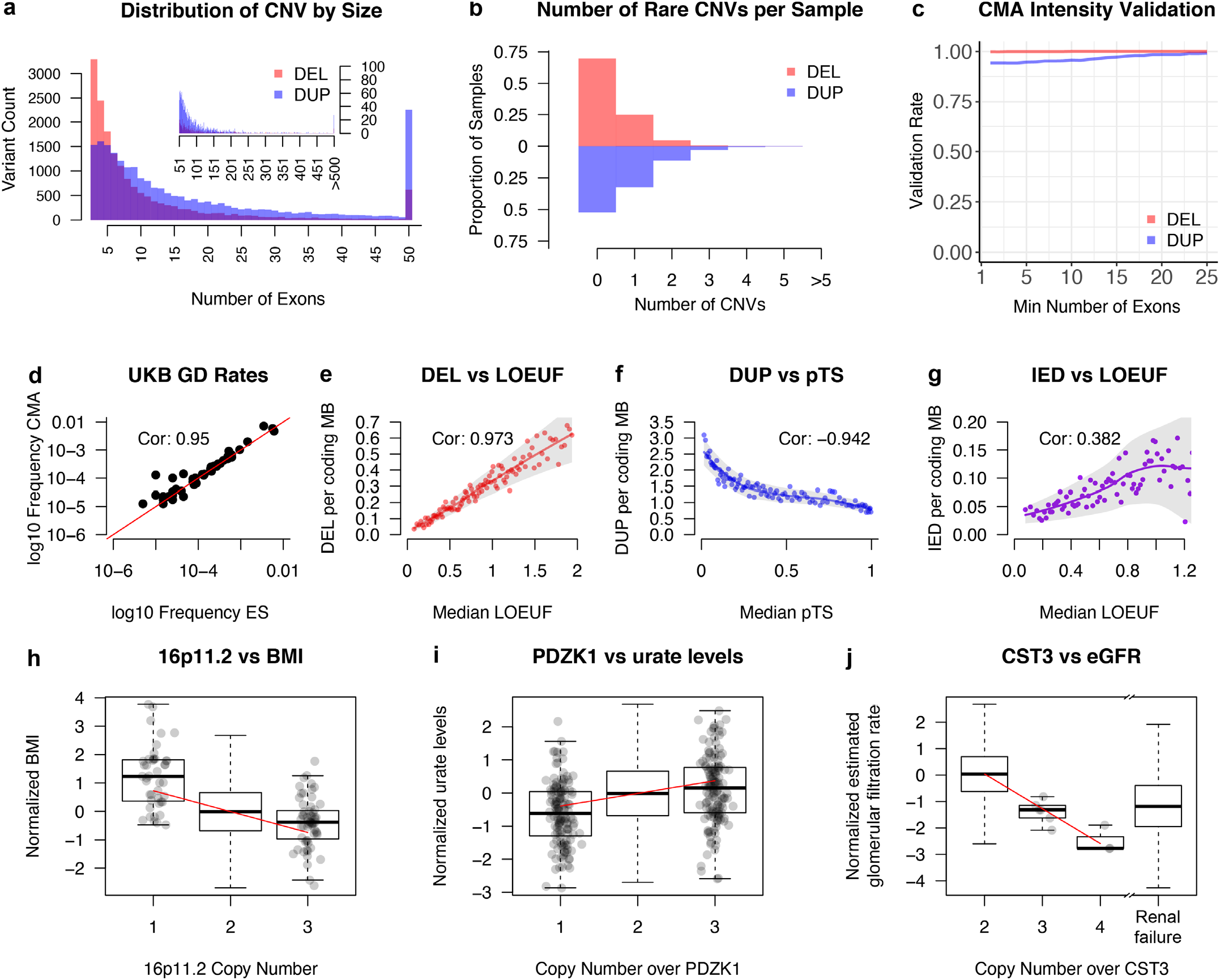
A high-quality rare CNV callset was generated on 200,624 exomes from the UK Biobank (UKBB) using GATK-gCNV. **a**, The variant-size distribution of high-quality, rare CNVs in the UKBB as a function of the number of exons each variant spans. **b**, The distribution of the number of rare, high-quality CNVs per-sample in the UKBB. **c**, Using 177,158 UKBB samples with matching CMA data, we find excellent validation of high-quality GATK-gCNV ES calls using Genome STRiP Intensity Rank Sum testing. **d**, GD CNV rates in the UKBB GATK-gCNV ES callset were highly concordant with rates from previous reports based on UKBB CMA data. **e**, The number of rare deletions observed over a gene in the UKBB GATK-gCNV callset is tightly correlated with LOEUF. **f**, The number of rare duplications observed over a gene in the UKBB GATK-gCNV callset is also strongly correlated with the pTS score measuring intolerance to duplications. **g**, The number of high-confidence duplications (IED) with both breakpoints within the boundaries of a gene are also correlated with LOEUF. **h**, 16p11.2 deletions are associated with a significant increase in normalized BMI. **i**, PDZK1 deletions are associated with a significant decrease in normalized urate levels. **j**, CST3 duplications are significantly associated with decreased normalized eGFR values, on par with eGFR of individuals with renal failure. **Abbreviations**: CNV - copy number variation; DEL - deletion; DUP - duplication; CMA - chromosomal microarray; UKBB - UK Biobank; LOEUF - loss-of-function observed over expected upper bound fraction; pTS - probability of triplosensitivity; IED - intragenic exonic duplication; GD - genomic disorder; ES - exome sequencing; BMI - body mass index; eGFR - estimated glomerular filtration rate.

In the absence of gold-standard GS data on all UKBB samples, we assessed the quality of the UKBB CNV callset generated by GATK-gCNV versus existing UKBB CMA datasets^32^. First, we conducted systematic in silico confirmation of high-quality variants from GATK-gCNV using the Intensity Rank Sum (IRS) test from the GenomeSTRiP software package^12^. We applied the IRS test to 33,679 high-quality sites from GATK-gCNV that (i) overlapped at least 10 CMA probes and (ii) exhibited site frequencies between 0.01% and 1% in the subset of 177,158 UKBB samples that had matching ES and CMA data available. For each variant, the IRS test compared the ordering of raw CMA probe intensities between predicted CNV carriers and non-carriers to determine if predicted CNV carriers have higher (for duplications) or lower (for deletions) CMA probe intensities, as would be expected for bona fide CNVs. This approach revealed that 95.7% of tested, high-quality CNVs from GATK-gCNV had orthogonal support from raw CMA intensity data at a nominal IRS p-value <0.01 (**Fig. 3c, Supplementary Fig. 5**). As a second, independent quality assessment of our ES-based UKBB CNV callset, we compared the rates of 49 genomic disorder (GD) CNVs^32^—large, disease-associated CNVs often formed by non-allelic homologous recombination—in our callset versus previously published rates from CMA analyses of the UKBB^32^. We found that the CNV frequency estimates at these 49 GD loci were highly concordant with prior CMA analyses of this same cohort (**Fig. 3d**, R^2^=0.95; P=1.5×10^−23^), providing further evidence for the accuracy of the GATK-gCNV-based UKBB CNV callset.

We next assessed whether the rates of CNVs in the UKBB correlated with established metrics of negative selection against LoF variation, or genic constraint. Several prior population-based studies, including those in the Exome Aggregation Consortium (ExAC) and the Genome Aggregation Database (gnomAD), have shown that negative selection against CNVs correlates with evolutionary constraint against LoF variation^4,26^, as measured by metrics like the LoF Observed over Expected Upper-bound Fraction (LOEUF^33^) from gnomAD or the probabilities of haploinsufficiency (pHI) and triplosensitivity (pTS) recently proposed by a large-scale CNV meta-analysis^34^. Encouragingly, we observed severe depletion of high-quality deletions in our GATK-gCNV callset that overlapped constrained genes as measured by both LOEUF (**Fig. 3e**) and pHI (**Supplementary Fig. 6**), as well as strong linear relationships between the number of deletions observed in UKBB per gene (defined as >10% deletion of exonic base pairs) and the constraint scores of those genes in percentiles (Spearman’s correlation=0.97 and =-0.97, respectively). Similarly, when examining the set of high-quality rare duplications from the GATK-gCNV callset, we find severe depletion in the number of CNVs that impact genes (defined as >75% duplication of exonic base pairs) found to be triplosensitive by pTS (**Fig. 3f**, Spearman’s correlation=-0.94), as well as a similarly strong linear relationship between the number of duplications and pHI (correlation=-0.97, **Supplementary Fig. 7**). Lastly, while the functional consequences of intragenic exonic duplications (IEDs) are context-specific and less readily predictable in silico, we nevertheless found depletion of putative IEDs correlating with genic constraint as measured by LOEUF (**Fig. 3g**, Spearman’s correlation=0.38), consistent with previous observations in gnomAD^26^.

Finally, as a demonstration of the utility of GATK-gCNV for trait association studies, we conducted a CNV-phenotype association analysis across 179,409 UKBB samples of European ancestry for a curated set of 478 traits (median: 177,400 samples per trait)^35^. We tested each phenotype for association against deletions and duplications of genes and against 46 previously reported GD loci^18^. After restricting to sites with at least 5 overlapping CNVs, we applied the Sequence Kernel Association Test (SKAT^36^) and adjusted for the top 20 SNP-based principal components^35^, sex, and age. At a conservative multiple-testing corrected threshold of 1.5×10^−8^ (**Supplementary Note**), we found 84 loci with significant associations (**Supplementary Table 1**), including a recapitulation of known GD-phenotype associations, such as the canonical 16p11.2 deletions with body mass index^37^ (BMI, p=1.9×10^−17^, **Fig. 3h, Supplementary Table 1**). Outside of known GD loci, we also identified associations in established pathogenic deletions, such as deletion of the hemoglobin gene cluster (encompassing *HBM, HBA2, HBA1, HBQ1*) which was previously associated with alpha thalassemia^38^. In our study we found this cluster to be significantly associated with the following alpha thalassemia-related blood traits^39^: mean corpuscular haemoglobin (p=3.5×10^−15^), mean corpuscular haemoglobin concentration (p=1.6×10^−12^), mean corpuscular volume (p=3.4×10^−15^), and erythrocyte count (p=2.4×10^−17^, **Supplementary Table 1**). We also recapitulated an association between deletions overlapping *PDZK1* and urate levels^40^ (**Fig. 3i**, p=1.6×10^−15^, **Supplementary Table 1**). Finally, we identified several potentially novel CNV-phenotype associations, with the most interesting involving a duplication of *CST3*, which was associated with an increase in cystatin C levels in blood (p=7.4×10^−17^) and a corresponding decrease in estimated glomerular filtration rate (eGFR, p=1.2×10^−21^). The decrease in eGFR tracked with increasing copy number (**Fig 3j, Supplementary Table 1**), providing support for the validity of this association. Curiously, we observed that individuals carrying *CST3* duplications presented with eGFR comparable to individuals in the UKBB with documented renal failure (n=5,455), although none of the *CST3* duplication carriers themselves were documented as having any renal-related disease phenotypes. Knowledge of these duplications could be clinically significant for these patients, sparing them the stress and follow-up testing for kidney diseases indicative of decreased eGFR levels.

## DISCUSSION

Despite the widespread usage of ES in clinical and research applications, the overwhelming majority of research studies using ES have not evaluated or leveraged CNVs and are limited to SNVs and indels alone. The advances in ES-based CNV discovery presented here using GATK-gCNV will provide significant added value to ES datasets. GATK-gCNV overcomes the challenge of read-depth signal normalization by implementing a flexible Bayesian model that removes known and unknown technical biases while preserving *bona fide* rare CNV signatures. We find that the recall and precision of GATK-gCNV to be suitable for use-cases ranging from association studies to sensitive diagnostic screening at a resolution of >2 exons when compared to gold-standard CNVs derived from GS. The critical feature of GATK-gCNV, which motivated its development, is its ability to maintain high accuracy for applications that require low false-positive rates, such as family-based research studies and clinical applications.

Despite the relative value added to standard ES applications, we highlight at least three significant limitations for GATK-gCNV studies. First, GATK-gCNV performance decays rapidly for CNVs smaller than three exons. Targeted individual exon analyses are routinely performed by visualization in many settings and single exon CNVs (in particular deletions) are often accessible with such an approach, but the sensitivity against GS in our analyses was insufficient for large-scale studies. Extracting read-depth data at a higher resolution and incorporating statistics beyond read-depth in the GATK-gCNV probabilistic model may improve the accuracy for smaller events in the future. Second, we have optimized GATK-gCNV for the detection of rare CNVs at a carrier frequency <1%; for common CNVs (frequency >1%), it becomes challenging to disentangle true polymorphic CNVs segregating in the general population from the technical biases introduced by probe-based hybridization capture. It is possible that these challenges may be mitigated in the future by incorporating prior weights on the distribution of population copy numbers at a given locus based on large, GS-based population databases of CNVs such as gnomAD^33^ or the UKBB^30^, and other approaches have been recently introduced that may accomplish this goal^41^. Third, all ES-based analyses are necessarily restricted in resolution to well-captured exons, and thus CNV breakpoint resolution is highly variable depending on local gene density. Although GATK-gCNV has provided utility for detecting rare exonic CNVs, ES data is fundamentally limited by the range of variation that is captured by targeted baits compared to the range of structural variants detectable genome-wide across the allele frequency spectrum from short-read or long-read GS.

This GATK-gCNV tool is fully accessible via the GATK software package (https://gatk.broadinstitute.org), where it can be deployed across local machines, high-performance enterprise computing clusters, and distributed cloud-computing environments (e.g., Google Cloud Platform, Amazon Web Services, Microsoft Azure). In addition, GATK-gCNV is fully supported via the GATK User Forum, which provides tutorials and example cloud workspaces. Using GATK-gCNV is relatively cost-efficient at $0.02-0.05 per ES sample, which could be further decreased through improved scaling using techniques such as amortized inference and subsampling. As a demonstration of the utility of GATK-gCNV, we applied it to 200,624 ES samples from the UKBB. These analyses serve to provide a resource of rare coding CNVs that we have released for use by the biomedical community (see **Data Availability**). We demonstrated patterns of CNV selection from GATK-gCNV wherein both coding deletions and intragenic duplications were depleted in genes constrained against LoF and that duplications were similarly depleted over genes predicted to be intolerant of increased gene dosage. As just one initial exploration of the myriad potential uses of this rare CNV resource in the UKBB, we demonstrate correlations between rare coding CNVs and several traits. We anticipate that the dissemination of these methods and data resources will catalyze new significant discoveries and deepen our understanding of the contribution of rare coding CNVs to a wide range of human traits and disorders.

## ACKNOWLEDGEMENTS

The authors would like to thank Lee Lichtenstein, Yossi Farjoun, Benjamin Neale, and Niall Lennon for insightful discussions at various stages of this project, and Shadi Zaheri for carefully reviewing and providing feedback on the manuscript.

This work was supported by grants from the Simons Foundation for Autism Research Initiative (#573206); the SPARK project and SPARK analysis projects (#606362 and #608540); the National Institutes of Health (MH115957, HD081256, HG008895, and HG011450). J.M.F. was supported by an Autism Speaks Postdoctoral Fellowship.

## AUTHOR CONTRIBUTIONS

M.B., D.I.B, and S.K.L. developed and implemented the GATK-gCNV model and the inference algorithm. A.S. contributed model enhancements and developed sample-clustering and batch-processing workflows. A.S., M.B, and S.K.L developed WDL workflows for Terra integration and scalable analysis. M.E.T, J.M.F., E.B, H.B., S.K.L., M.W., and L.D.G. supervised aspects of this project at various stages of development. J.M.F., R.L.C., H.B., and K.J.K. contributed to association analyses. I.W. and J.M.F. generated the CNV callsets. J.M.F., I.W., R.L.C., A.S.-J., and H.B. conducted quality-control on generated callsets.

## CODE AND RESOURCE AVAILABILITY

### Code Availability

GATK-gCNV is distributed as part of the GATK jar release. For an example workspace on Terra, with recommended parameters, please see:

https://app.terra.bio/#workspaces/help-gatk/Germline-CNVs-GATK4

### Data Availability

The UKBB CNV dataset has been returned to the UKBB for distribution to approved researchers.

## Online Methods

### Overview of GATK-gCNV probabilistic approach to CNV detection

GATK-gCNV employs a generative model of sequencing read-depth data that accounts for both a) copy number variation and b) technical variation associated with differences in sample extraction, library preparation, enrichment, sequencing, and mapping. The method takes as input the read-depth data from a collection of samples over a set of genomic intervals, and learns to disentangle CNV events from technical factors by modeling both on an equal footing. Conceptually, disentanglement is made possible due to the discreteness and rarity of germline CNV events relative to the continuity and ubiquity of technical variation. Our proposed generative model consists of two main compartments, a model for read-depth likelihood given the copy number states, and a hierarchical hidden Markov model (HHMM) that encodes copy number prior probabilities and state transitions along the genome. The read-depth likelihood compartment is a negative-binomial linear latent-factor model that accounts for technical read-depth variation in terms of a small number of learnable and predetermined bias factors over the genomic intervals. Individual samples share statistical power by determining the shared bias factors together. The copy number state HHMM compartment models the copy number structure, both at the level of individual samples and at the population level, and accounts for the genomic correlation of copy number states and the higher state-to-state transition rate within CNV loci that are determined to be polymorphic. Model parameters and latent variables, including copy number states and read-depth bias factors, are inferred simultaneously within a variational inference framework. We will describe the GATK-gCNV model and inference in the following sections. Further technical details, in particular those pertaining to implementation and the inference algorithm, are provided in the **Supplementary Note**.

### Likelihood model for read depth conditioned on copy number state

We consider the integer read-depth matrix *n*_*st*_, with rows *s*= 1,2, …,*S* and columns *t*=1,2, …,*T* denoting samples and genomic intervals, respectively (**Supplementary Fig. 1a**). Our goal is to model the conditional distribution *p*(*n*_*st* |_ *c*_*st*_), where *c*_*st*_ is an integer copy number state matrix. At a fundamental level, the observed counts are typically obtained by sequencing a random subsample of the short-read hybrid capture-based library. As such, we expect random sampling noise (i.e., Poisson noise) to set the lower bound on the count dispersion. In practice, this fundamental noise is far outweighed by other sources of systematic noise, such as amplification artifacts and sequencing biases that are difficult to explicitly model. We take a data-driven approach and model *n*_*st*_ as a negative-binomial (NB) distributed random variable with λ_*st*_ ≥ 0 rate and overdispersion Ф_*st*_ ≥ 0:

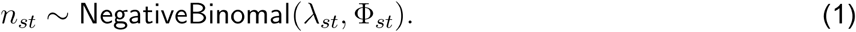

In our choice of NB parameterization, 𝔼 [*n*_*st*_] = λ_*st*_ and 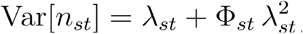. Our general approach to modeling is to capture the generalizable patterns of read-depth variation in the NB rate λ_*st*_, and to allow the NB overdispersion to absorb the residual variance. To this end, we structure the NB rate λ_*st*_ into multiplicative contributions arising from sequencing depth, copy number, and capture bias, as well as a small additive contribution from read-mapping errors:

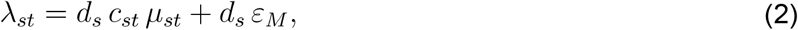

where *d*_*s*_ ∼ Log Normal (*μ*_*d*_, σ_*d*_) is the sample-specific sequencing depth with prior mean *μ*_*d*_ and standard deviation σ_*d*_ as model hyperparameters, is the integer copy number matrix, ε*M* is a small mapping-error rate hyperparameter, and *μ*_*st*_ is the multiplicative bias factor matrix. We model the latter as a low-rank linear latent-factor model with an exponential link function:

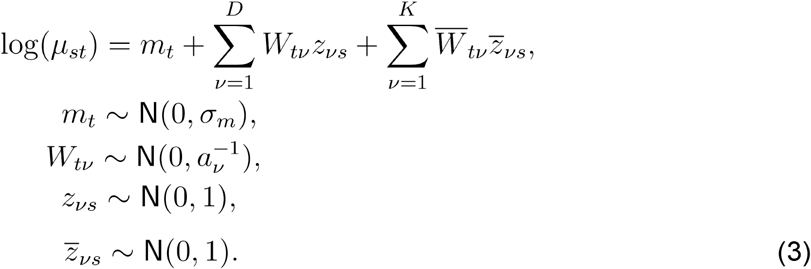

Our model for the bias matrix *μ*_*st*_ comprises three terms: (I) The first term is a target-specific mean bias *m*_*t*_ that is shared across all samples and has a normal prior with scale σ_*m*_ as a model hyperparameter; (II) The second term is a product of *D* learnable bias factors 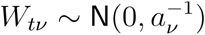 and their corresponding sample-specific loadings *𝓏*_*vs*_ ∼N (0,1). This structure can be thought of as a factor analysis sub-model. During model-fitting, all samples contribute to learning the same bias factors *W*_*tv*_, whereas each sample uses (“loads”) the factors to varying degrees. We set the number of bias factors *D* to an estimated upper bound, e.g. *D* ∼ 10 − 20, and tune the prior scale of each factor 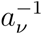 to maximize model evidence. Known as automatic relevance determination (ARD), this empirical Bayes procedure shrinks the prior scale of unnecessary bias factors to zero and automatically selects the appropriate number of bias factors from the data. (III) Finally, the last term is a product of *K* predetermined bias factors 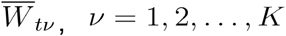 and their corresponding sample-specific loadings 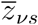. This provision allows us to explicitly include known read-depth bias factors into the model and accelerate model training. In practice, we found it beneficial to treat the GC-content of target genomic intervals as predetermined bias factors. To this end, we set a lower and upper bound on the GC-content according to our interval filtering criteria and binned the allowed range uniformly into equally-sized bins. We determined the GC-content of each genomic interval as a preprocessing step, constructed a mapping φGC : *t* → 1, ⋯,N_GC_ from each genomic interval to the best-matching GC-content bin, and set the GC bias factors as, 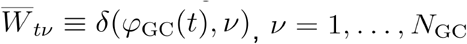. Intuitively, 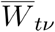 selects all genomic intervals with similar GC contents and the inferred sample-specific loadings 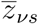 can be thought of as the conventional “GC curves”. We did not include any other hand-crafted bias factors in our implementation and therefore,*K* = *N*_*GC*_.

Finally, we allow the likelihood model to capture the variance that is not accounted for by the described bias-factor model using the NB overdispersion Φ_*st*_. We propose the following parametric decomposition of the overdispersion into target-specific and sample-specific contributions:

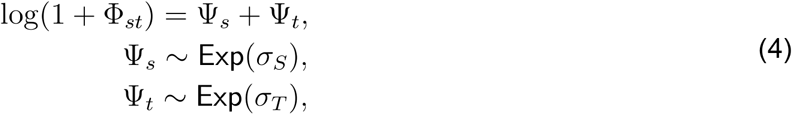

where σ*S* and σ*T* are model hyperparameters. The NB overdispersion can be thought of as a stopgap mechanism to prevent overfitting. Without this mechanism, model misspecification will lead either to learning non-generalizable bias factors, or worse, exploitation of the copy number state variables *c*_*st*_ as well as the genomic-region class *τ*_*t*_ latent variables (defined below) to account for the residual variance. It is easy to show that Eq. (5) induces a heavy-tailed distribution over Φ_*st*_. This permissive prior allows the bias latent-factor model to “fail fast,” effectively preventing overfitting and ultimately increasing the precision of the detected CNVs.

### Hierarchical Hidden Markov Model for copy number states

We model the copy number state prior probabilities via a two-level hierarchical hidden Markov model (HHMM) as shown in **Supplementary Fig. 1b**. The top-level, primary Markov chain dictates the “class” of a genomic region as active (highly polymorphic) and/or silent (mostly diploid); this binary determination, in turn, sets the prior probability and the state-to-state transition matrix of the secondary Markov chains. Active regions are given permissive copy number priors (i.e. uniform, **Supplementary Fig. 1c**), while silent regions have priors heavily weighted on the copy-neutral state (**Supplementary Fig. 1d**). Adjacent genomic regions are more likely to belong to the same region class, and we model this using a “sticky” region-to-region transition matrix.

Next in the hierarchy is a group of Markov chains, one for each sample, and conditionally independent of one another given top-level region-class variables. These secondary chains model the state-to-state copy number transitions along the genome separately for each sample. Again, genomic regions within a characteristic length scale tend to have similar copy number states, which we also model using a “sticky” copy number state-to-state transition matrix.

We describe both levels of the hierarchy in more detail in the following sections.

### Top-level Markov chain: genomic-region classes

To model highly polymorphic (active) and mostly diploid (silent) genomic regions in a unified model, we introduce a per-interval binary random variable τ_*t*_ ∈{active,silent} (*t* = 1, ⋯,*T*) . The region class of the first interval is sampled from a Bernoulli distribution, τ_1_ ∼ Bernoulli (π_region_), where:

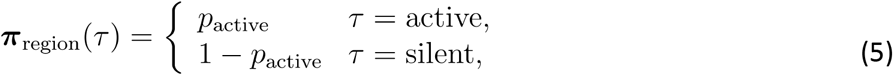

where *P*active is a model hyperparameter. The region classes of subsequent loci are conditionally sampled according to the following transition matrix:

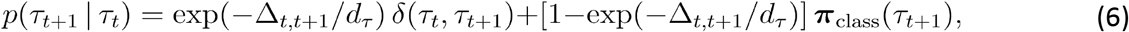

where Δ*t,t* + 1 is the genomic distance between the midpoints of region *t* and *t* +1, *d*_*τ*_ is a model hyperparameter that determines the typical correlation length of region classes, and is the Kronecker delta function. Eq. (7) models the “sticky” behavior advertised earlier and is best understood by considering two limiting cases: (1) in the limit Δ*t,t* + 1 ≪ *d*_*τ*_, we obtain *p*(τ_*t*+1_| τ_*t*_) ≈ δ (τ_*t*_, τ_*t*+1_), i.e. the next region inherits the state of the previous region; (2) in the limit Δ*t,t* + 1 ≪ *d*_*τ*_, we obtain *p*(τ_*t*+1_| τ_*t*_) ≈ π _*τ*_ (τ_*t*+1_), i.e. the previous state is forgotten and the next region is sampled from the prior.

### Secondary Markov chains: sample-specific copy number states

Given a determination of the genomic-region classes from the top-level chain, the copy number states of each sample (i.e. the rows of the copy number matrix, see **Supplementary Fig. 1**) are independent of one another. We set an upper bound on the largest detectable copy number,*C*, as a model hyperparameter. We further assume being given a matrix of baseline copy number states for each sample and at each genomic region, *𝓀*_*st*_. For a diploid organism, *𝓀*_*st*_ =2 in the autosome (except for samples with aneuploidy) and *𝓀*_*st*_ = 0,1, 2 for sex chromosomes (depending on the per-sample sex-chromosome ploidy). We interpret the copy number matrix *c*_*st*_ as a small perturbation of the baseline copy number matrix *𝓀*_*st*_. We define the prior copy number distributions for the silent and active classes as follows:

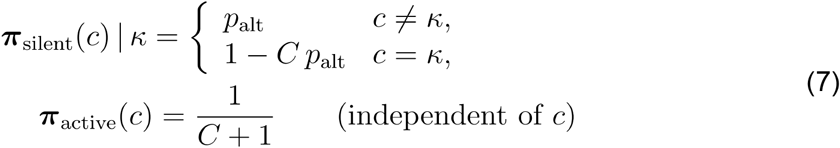

These priors are schematically shown in **Supplementary Fig. 1b**. Note that is another model hyperparameter that determines the permissiveness of having a non-baseline (e.g., non-diploid) copy number state in silent regions. The prior distribution is assumed to be flat in active regions, that is, all *C* + 1 copy number states are assumed to be equally likely.

At the first interval *t* = 1, the copy number state in sample *s* is sampled from the prior:

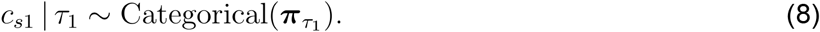

For the subsequent targets, the copy number state is sampled according to the following transition matrix:

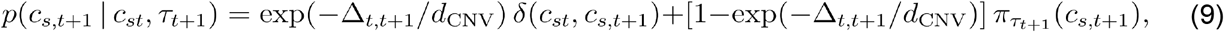

where Δ*t,t* + 1 is the genomic distance between the midpoints of region *t* and *t* + 1 as before, and is a model hyperparameter that determines the typical correlation length of CNV events. Eq. (10) models the “sticky” behavior advertised earlier and is again best understood by considering two limiting cases: (I) in the limit Δ*t,t* + 1 ≪*d*_CNV_, we obtain *p*(*c*_*s*_,_*t*+1_ | *c*_*st*_, τ_*t*+1_) ≈ δ(*c*_*st*_,*c*_*s*_,_*t*+1_), i.e. the next region inherits the copy number state of the previous region; (II) in the limit Δ*t,t* + 1 ≫*d*_CNV_,, we *p*(τ_*t*+1_| τ_*t*_) ≈ δ (τ_*t*_, τ_*t*+1_), obtain *p*(*c*_*s*_,_*t*+1_ | *c*_*st*_, τ_*t*+1_) ≈ π _τ_ (τ_*t*+1_), i.e. the previous state is forgotten and the next region is sampled from the prior.

### Determining chromosomal baseline copy number states

The generative model for copy number states requires the knowledge of the chromosomal baseline copy number matrix 𝓀_*st*_ for each sample *s* = 1, ⋯*S* at genomic interval *t* = 1, ⋯,*T*. By definition, the baseline copy number is the most prevalent copy number state at the scale of chromosomes (e.g., 2 for diploid, 3 for trisomy, etc.), and its determination serves to unify the treatment of diploid and aneuploid samples, as well as sex chromosomes in mixed-sex sample cohorts. All genomic regions belonging to the same chromosome *j* = 1, ⋯,*J* have the same baseline copy number and therefore, it is sufficient to determine a copy number matrix, 𝓀_*sj*_, at the resolution of chromosomes instead of fine-grained genomic targets. We define the per-chromosome read-depth as:

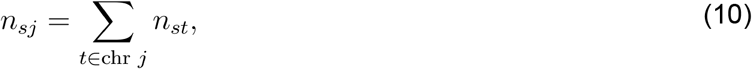

and like before, model it as negative-binomial distribution:

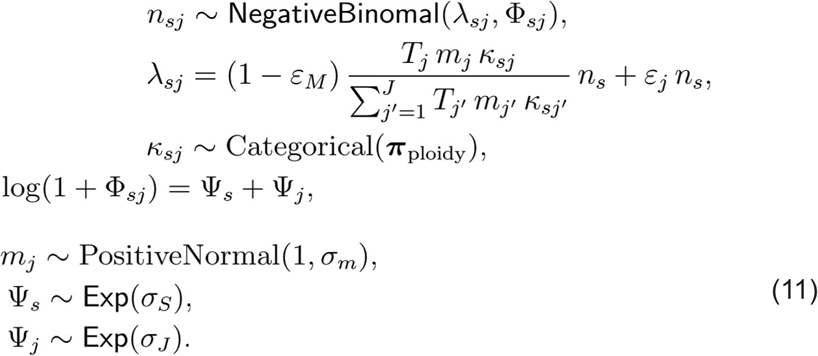

Here, *T*_*j*_ = | {*t* : *t* ∈ chr *j*} is the number of genomic intervals spanning chromosome *j*, ε *M* is a mapping error rate,*n*_*s*_ = ∑_*t*_ *n*_*st*_ is the sample-wide total read-depth, and 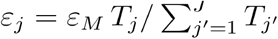 is the fractional mapping error rate for chromosome *j*. The multiplicative bias *m*_*j*_ accounts for chromosome-to-chromosome bias in read-depth. Like the fine-grained read-depth model, we account for the unexplained chromosome-scale read-depth variance as a sum of sample-specific Ψ*s* and chromosome-specific Ψ*j* contributions. Finally, π_ploidy_ is the chromosome-scale ploidy prior.

### GATK-gCNV model fitting using variational inference

The structured Bayesian model we described above captures key aspects of the phenomenology of sequencing read-depth variation and germline CNV events in a unified manner. However, the complexity of this hierarchical model and the lack of simplifying Bayesian conjugacy relationships implies that an exact inference algorithm is likely to be out of reach. Practical approximate-inference strategies include sampling-based Markov chain Monte Carlo (MCMC) methods and variational inference (VI). Here, we pursue VI as a more attractive option for the following reasons: (I) VI typically allows faster convergence times compared to MCMC-based strategies; (II) the flexibility of VI allows us to perform exact inference on certain sectors of the model (i.e., copy number HMMs); (III) recent advances in machine-learning software and probabilistic programming languages (PPLs) allow us to perform automated VI over the continuous sector of the model (i.e., the read-depth likelihood compartment) with little effort. We describe the details of our variational-inference approach in **Supplementary Note**. Operationally, we adopt a mean-field approximation and neglect posterior correlations between continuous latent variables **Z**_continuous_ (e.g., sequencing depth, bias factors, loadings, etc.) and discrete latent variables **Z**_discrete_(e.g., copy number states and genomic-region class indicators). We further assume a fully-factorized mean-field posterior for **Z**_continuous_ and neglect posterior correlations between top-level and secondary Markov chains in the HHMM compartment. We leverage the PyMC3^41^ PPL to perform incremental variational updates of the continuous posterior. These updates are interleaved with updates of the discrete posterior distributions, which are made tractable by exploiting the emergent linear conditional random field (CRF) structure that follows from mean-field factorization. An annealed entropy-regularization strategy is used throughout to avoid poor local minima in the early stages of model fitting, and convergence is assessed by testing the stability and self-consistency of variational posteriors within specified error tolerances.

### Code accessibility and resource availability

https://app.terra.bio/#workspaces/help-gatk/Germline-CNVs-GATK4

## Supplementary Note

### A hybrid auto-differentiation variational inference (ADVI) framework for GATK-gCNV

Consider a model with a bundle of tunable parameters Θ, latent variables **Z**, and observed data **X**. In theory, one wishes to obtain an optimal parameterization of the model by maximizing the model evidence, *p*(X | Θ) (also known as the marginal likelihood). An exact calculation of the model evidence, though, requires marginalizing the latent variables **Z** and is practically out of reach. Variational inference (VI) relies on optimizing a lower bound of the model evidence (ELBO) as obtained via Jensen’s inequality:

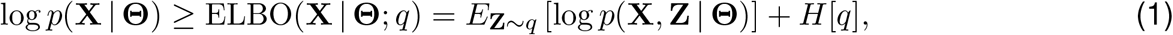

where *q*(Z | **φ**) is a *variational distribution*, parameterized by **φ** and aiming to approximate the true posterior *p*(**Z** | **X, Θ**) as closely as possible, and *H*[*q*]= −*E*_***Z***∼*q*_ [log *q*(**Z** | **φ**)] is the entropy of *q*(*Z* | **φ**). The objective of VI is to maximize ELBO over both model parameters **Θ** and posterior variational parameters ***‘*** and thereby reducing the gap to the actual model evidence.

For Bayesian models with fully continuous bundle of latent variables **Z**, an effective automated strategy is (1) choosing a simple variational form, e.g. fully or partially factorized Gaussian distributions with their respective means and variances as variational parameters φ, (2) approximating the **Z** expectations appearing in the ELBO (Eq. 1) with a finite number of MC samples, and (3) leveraging autodifferentiation tensor computation frameworks (e.g. Theano, PyTorch, or TensorFlow) to automate the calculation ∇ _***φ***_ ELBO(**X** | **Θ**; *q*), and (3) optimizing the parameters via gradient descent updates. This procedure is termed auto-differentiation variational inference (ADVI) and is implemented in several probabilistic programming packages such as PyMC3, Pyro, and TensorFlow Probability (TPF). ADVI and its extensions unburden model developers from hand-deriving expectation-maximization and variational Bayes update equations, and enables carrying out model criticism and model update in faster iterations. The ADVI framework, however, is not readily applicable for models that involve discrete latent variables or a mixture of discrete and continuous latent variables. In practice, the variance resulting from approximate marginalization of discrete latent variables (via finite MC samples) is too high to allow fruitful gradient updates. Devising general-purpose and automated inference strategies in such situations is an active area of research in the probabilistic programming field.

A high-level inspection of our generative model provides clues for devising a VI scheme that is suitable for the present model. On the one hand, each of the HMM chains appearing in the copy-number matrix submodel, conditional on the top-level chain, admits a closed-from exact posterior via the forward-backward message passing. On the other hand, the discrete and continuous latent variables appear, by and large, in separate cliques. For instance, the *only* discrete latent variable appearing in the read-depth likelihood submodel. We can exploit the existence of these model features to devise a *hybrid* VI strategy that leverages ADVI for continuous variables and message passing for discrete latent variables. We refer to this strategy as *Hybrid ADVI* (to be outlined below in generality), and we will more concretely discuss the resulting parameter update steps in the context our model in the next sections.

Let us define **Z** = **Z**_D_ ∪ **Z**_C_, where **Z**_D_ and **Z**_C_ are mutually exclusive discrete and continuous latent variables, respectively. Without the loss of generality, we can decompose the log joint probability as follows:

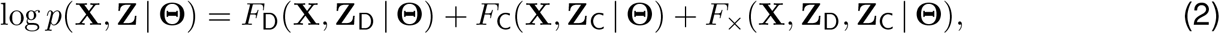

where *F*_D_ and *F*_C_ only depend on Z_D_ and Z_C_, respectively, and *F*_×_ involves both. A canonical choice is to set *F*_D_ and *F*_C_ to be the largest discrete and continuous cliques, in the absence of cross terms, and bundling the remaining potentials as *F*_×_. At this point, we make a number of simplifying assumptions.

First, in the spirit of ADVI which assumes fully-factorized posteriors, we consider the class of variational posteriors that are at least factorized in discrete and continuous sectors:

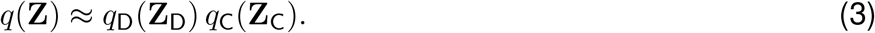

We will impose additional (approximate) factorization assumptions separately over *q*_D_ and *q*_C_ later on. For the time being, the high-level discrete vs. continuous factorization is sufficient. Second, we assume that none of the model parameters Θ appears in the cross term *F*_×_ (or otherwise, their contribution is negligible):

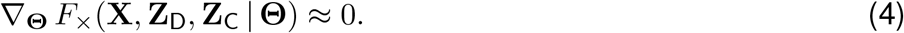

After plugging Eqs. (2) and (3) into Eq. (1) and making several straightforward rearrangements, we find:

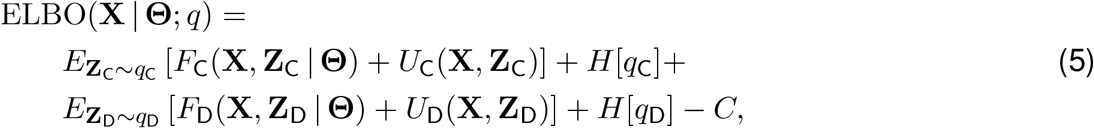

where:

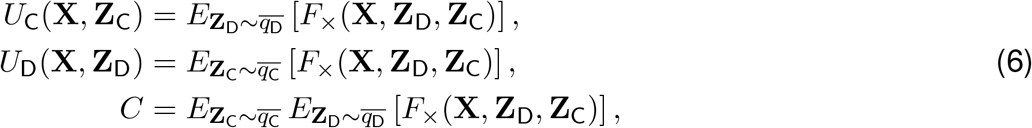

where we have used the following identity:

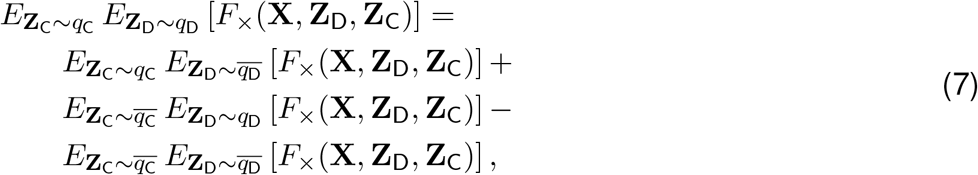

where 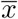 implies the *stop-gradient operator*, meaning the expression is to be treated as a constant in gradient calculations. We note that (1) Eq. (7) is only valid for evaluating the expression and its first derivatives, though, not for higher order derivatives (which are not needed here); (2) the constant term *C* can be dropped from Eq. (5) as both variational posteriors have stop-gradient and the cross potential has no Θdependence. The decomposition of the ELBO obtained in Eq. (5) has an intuitive mean-field interpretation. The discrete and continuous random variables are endowed with their ELBO-like optimization objectives in which the respective mean-field cross potentials, *U*_D_ and *U*_C_, have stop-gradient over the opposite sectors, *q*_C_ and *q*_D_. In practice, this partitioning leads to interleaved and independent updates of *q*_C_ and *q*_D_, each increasing the overall ELBO. The continuous ELBO can be re-interpreted as a Bayesian model that is free of discrete random variables and thus, can be optimized using ADVI. The discrete ELBO, after mean-field decoupling of top-level and copy-number chains, can be identified as the ELBO of independent linear-chain conditional random field (CRF) models and thus, can be treated via exact message passing without the need to resort to further mean-field assumptions.

### The continuous sector: auto-differentiation variation inference (ADVI)

We recall that for the present model, X = {*n*_*st*_} is the read-depth matrix, Θ comprises the ARD precision coefficients {*a*_ν_ : ν = 1,. .., *D*}, Z_D_ comprises all latent variables except for {*c*_*st*_} and {τ_t_}, and **Z**_C_ comprises only {c_st_} and {τ_t_}. We recall that the effective ELBO of the continuous sector of the model is given as:

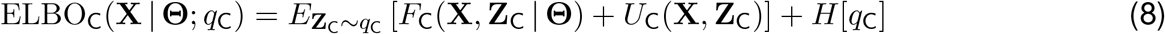

where:

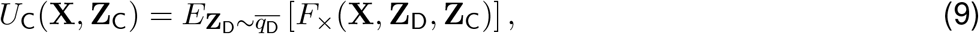

is the effective mean-field potential that arises from coupling of discrete variables to continuous latent variables.

A straightforward inspection of our model reveals that the only potential that contains cross terms of discrete and continuous variables is the clique associated to *n*_*st*_ | *c*_*st*_. The associated potential of this clique is:

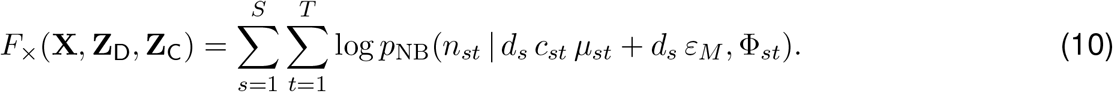

Given our current belief of the posterior distribution of copy-number states *q*(*c*_*st*_), the effective potential for the continuous ELBO, *U*_C_, is therefore simply given by the following local summation:

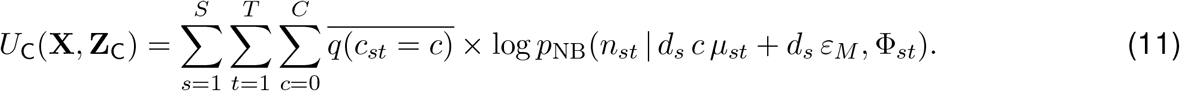

In practical terms, performing ADVI over Z_C_ is accomplished by replacing the negative binomial log probability with the effective potential given above.

### The discrete sector: variational message passing

Similarly to the previous section, the effective ELBO of the discrete sector of the model is given as:

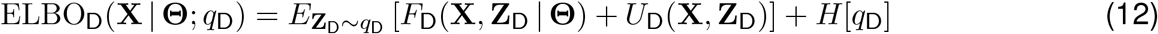

where:

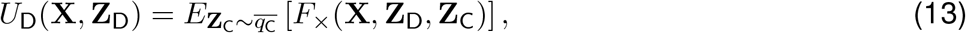

is the effective mean-field potential that arises from coupling of continuous latent variables to discrete latent variables. In the context of the present model, the cross potential *F*_×_ is given by Eq. (10) and only involves *c*_*st*_. Furthermore, the full cross potential is obtained as a sum of potentials that only depend on individual entries of the copy-number matrix *c*_*st*_. More explicitly:

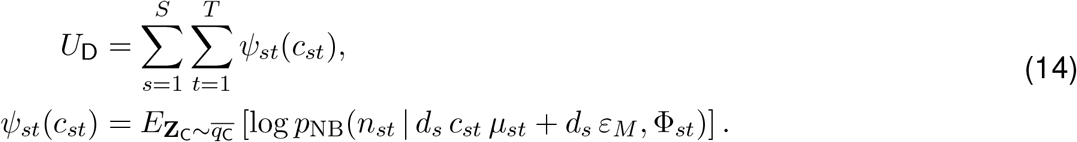

In practice, we estimate *ψ*_*st*_(*c*_*st*_) via a finite number of Monte Carlo samples, *n*_MC_, from our current belief of the posterior of continuous variables, *q*_*C*_. We refer to {*ψ*_*st*_} as the *effective emission potentials*, in accord with the HMM terminology.

Performing an exact update of *q*_*D*_ is conceivable, however, is computationally intractable. It is straightforward to show that optimizing ELBO_D_ amounts to performing inference on a HMM with a state space of size 2 × (*C* + 1)^*S*^; factor of 2 for the top-level chain, and a factor of *C* +1 for each of the per-sample copy-number states; see **Supplementary Fig. 1c**. As an approximate alternative, we resort to the following argument: for *S* ≫ 1, we expect q_*τ*_ to become exceedingly sharp as many samples contribute to determining the binary genomic re_T_gion class, active or silent. In the *S* → ∞ limit, the region class will be fully determined, i.e. 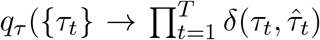, where 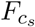 is the actual region class. The determinism of {*τ*_*t*_}, in turn, leads to decoupling of copy-number Markov chains of different samples from one another. In other words, the following factorization of the discrete posterior,

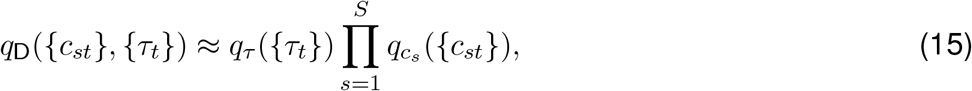

becomes increasingly more accurate as the cohort size *S* increases, and becomes exact in the limit *S* → ∞. Assuming finite but large *S* in practice, we can use such a mean-field factorization as a descent variational ansatz. The The expression for *F*_D_ can be spelled out from **Supplementary Fig. 1c**:

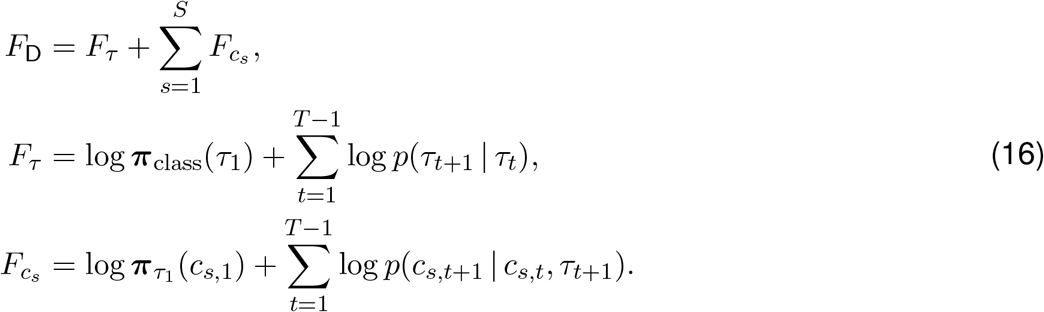

The linear-chain structure of both *F*_*τ*_ and 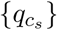, together with the mean-field ansatz Eq. (15), imply that the effective ELBO obtained for either {*τ*_*t*_} or {*c*_*s,t*_} is identical to the ELBO of a generic linear-chain conditional random field (CRF) with the following factor graph:

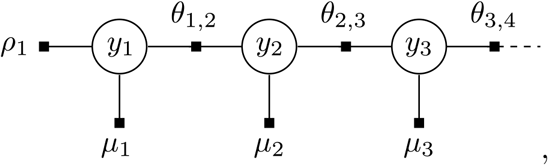

where *y*_*t*_ is either *τ*_*t*_ or *c*_*st*_, along with appropriate choices of the factors *ρ*_1_, {*µ*_*t*_}, and {*θ*_*t,t*+1_}. The probably distribution corresponding to such a linear-chain CRF is formally given as:

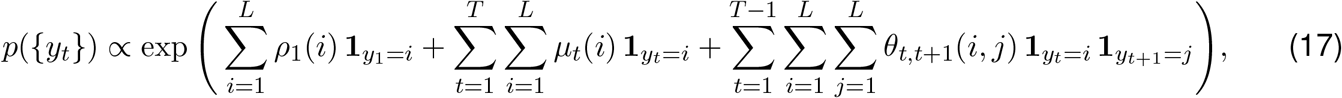

where *L* is the maximum number of states, i.e. *L* = 2 and *L* = *C* +1 for region class and copy-number linear-chains, respectively. Crucially, local marginals, i.e. *p*(*y*_*t*_), and marginals of adjacent pairs of nodes, i.e. p(*y*_*t*-1_, *y*_*t*_), can be obtained efficiently using the forward-backward message passing algorithm (see Algorithm 1). The factors can be readily worked out using Eqs. (12), (14), (15), and (16). We provide the expressions here for reference. For each of the S copy-number chains, we obtain the following factors:

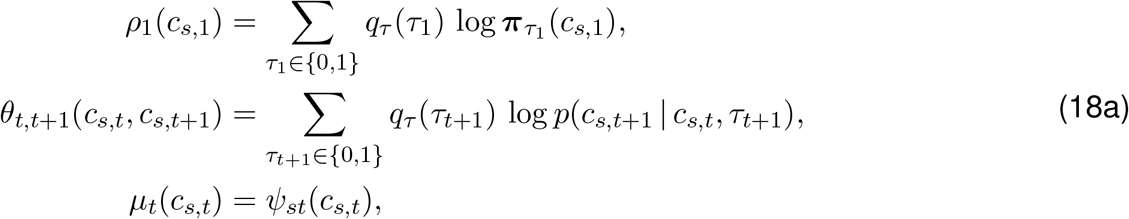

and for the region class chain, we have:

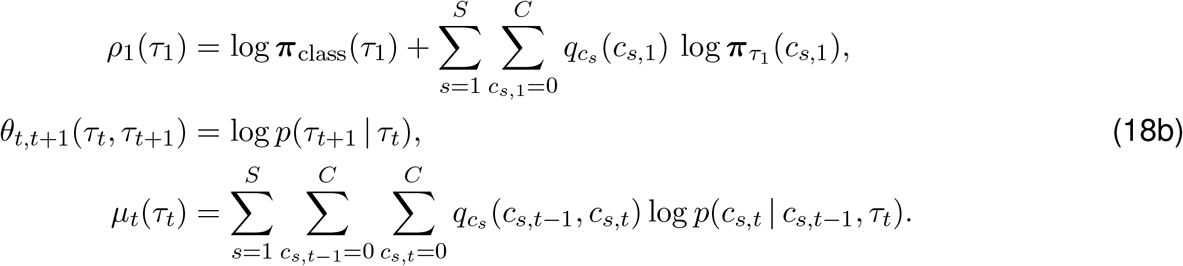

Note that the factors of the *τ*-chain depends on marginals of the other *c*_*s*_-chains, and vice versa, as expected from the factorized variational posterior we have used (Eq. 15). A self-consistent solution for *q*_*τ*_ and 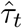 can be obtained via interleaved updates and relaxation. The detailed algorithm is given in Algorithm 2. In practice, we achieve convergence within < 5 interleaved updates using an admixing (relaxation) coefficient *α* = 0.75.

#### Algorithm 1 Forward-backward message passing for linear-chain CRF

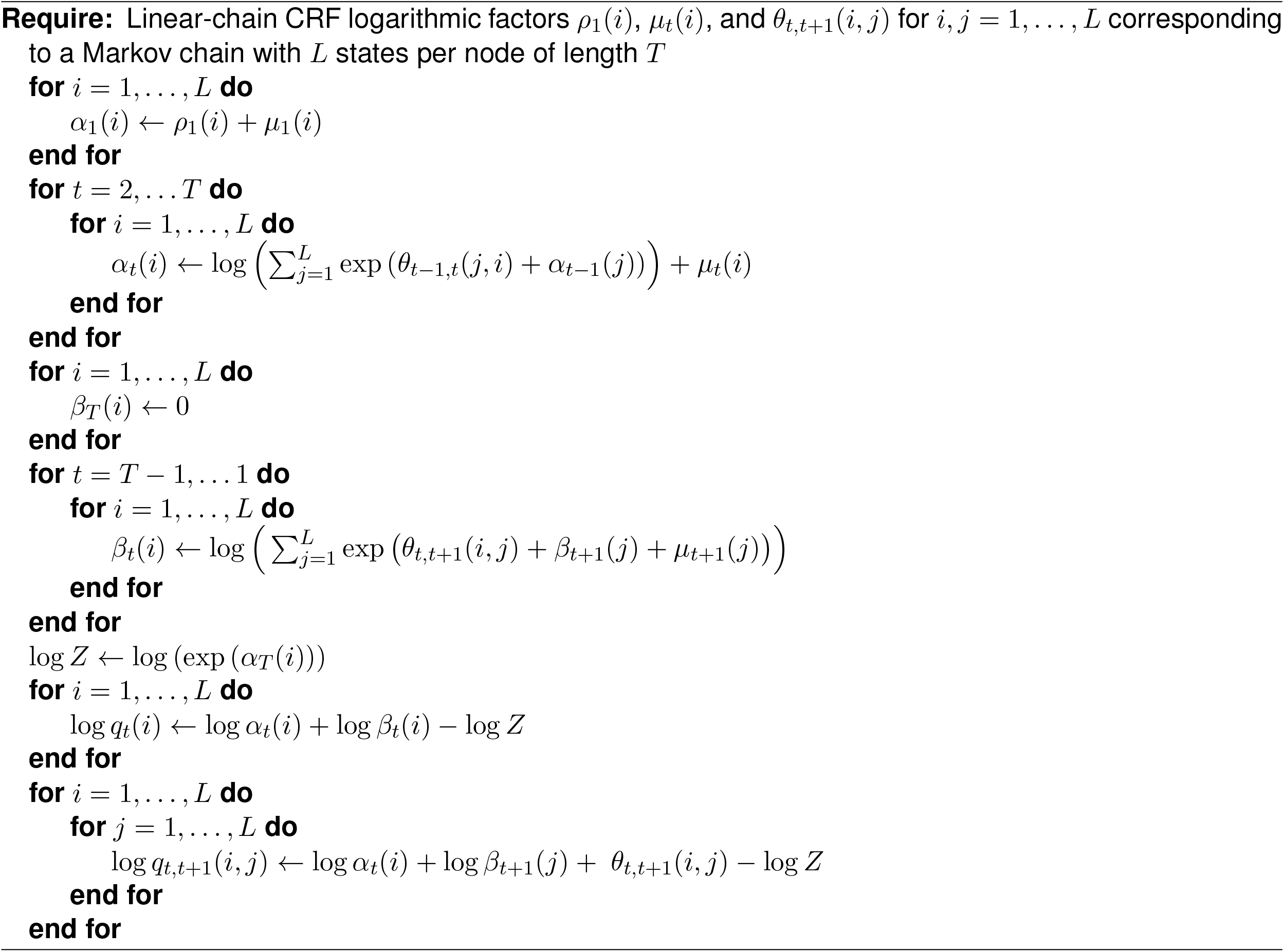

#### Algorithm 2 Posterior update of the discrete latent variables of the GATK-gCNV model

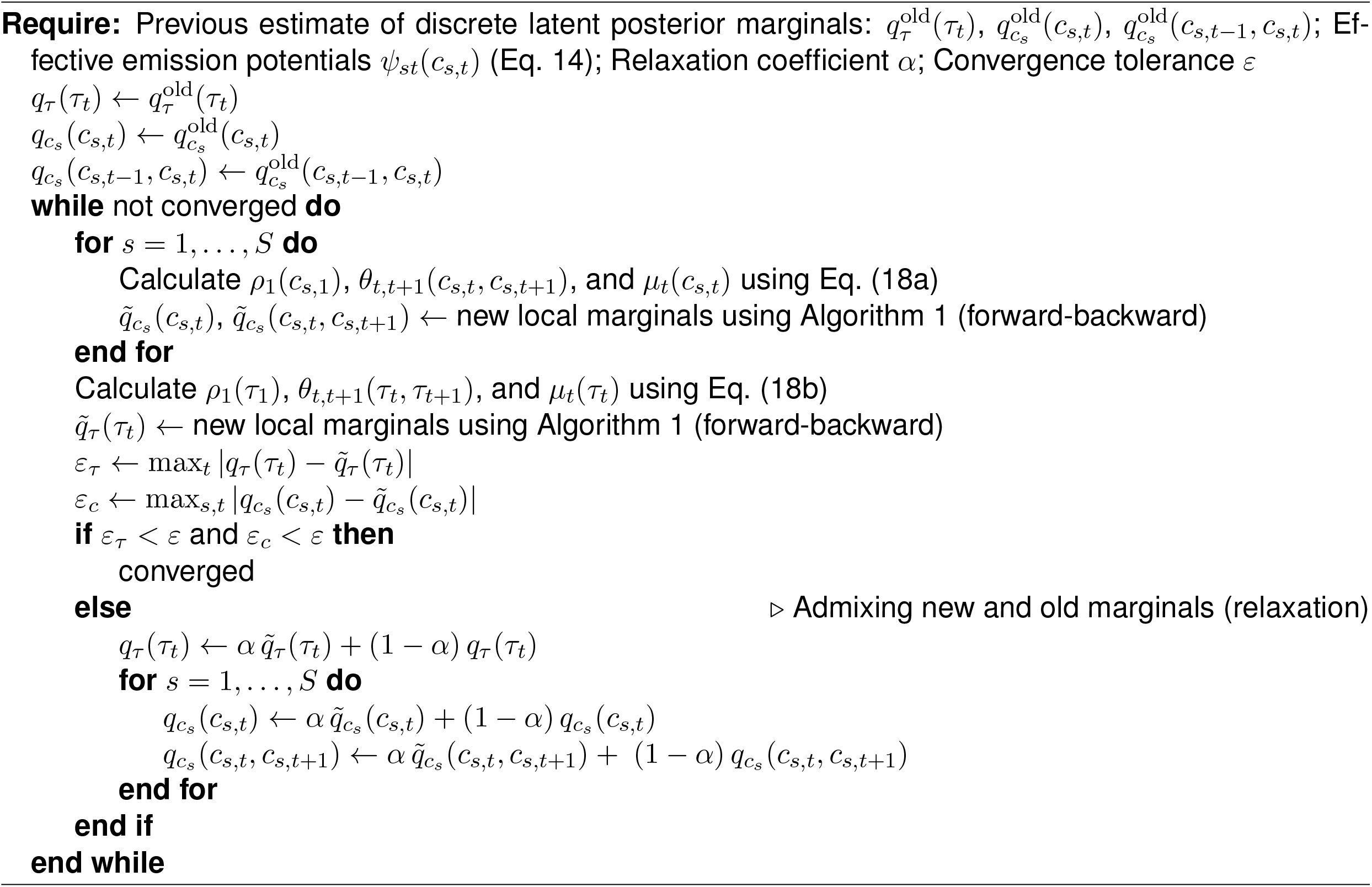

### Hybrid ADVI for the baseline copy-number state model

The Hybrid ADVI inference algorithm for inferring baseline copy-number states according to the model shown in **Supplementary Fig. 1b** is similar to the interleaved inference algorithm described in the last two sections. In this rather simpler model, Z_D_ = {𝒦_*sj*_}, and Z_C_ = {*m*_*j*_, Ψ_*s*_, Ψ_*j*_}. The cross potential arises from the clique *n*_*sj*_ | 𝒦_*sj*_, and the associated expression is:

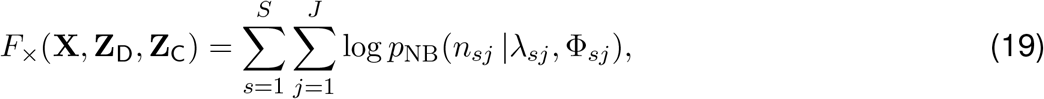

where the explicit expression for λ_*sj*_ is given in Eq. (11) in **Online Methods**. To perform ADVI update over Z_C_, we marginalize 𝒦_*sj*_ given the current posterior belief of baseline copy-number states *q*_𝒦_ (𝒦_*sj*_). To perform update of the latter, we calculate the effective emission potential via Monte Carlo sampling:

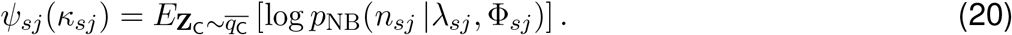

The update equation for *q*_𝒦_ is found by maximizing ELBO_D_, which coincides with the Bayes rule:

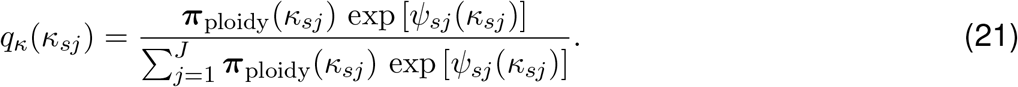

The ADVI update of *q*_C_({*m*_*j*_, Ψ_*s*_, Ψ_*j*_}) and Bayes rule update of q_𝒦_(𝒦_*sj*_) are continued in an interleaved fashion until a self-consistent and stationary solution is obtained.

### Implementation details

A detailed account of the GATK-gCNV variational inference algorithm was given in the previous two sections. We conclude our exposition here with a brief account of the implementation details. We have implemented the GATK-gCNV probabilistic model and inference algorithm as an encapsulated Python module named `gcnvkernel`, as a part of the GATK code base. We extensively leverage `Theano` tensor computation system and `PyMC3` probabilistic programming language in `gcnvkernel`. The tool can operate in two different modes: *cohort* mode, and *case* mode. In the cohort mode, the gCNV model is fit to a collection of samples and the tool outputs (1) a trained model, and (2) inferred copy-number states and various quality metrics (to be described later) for each sample in the cohort. In the case mode, the tool consumes global latent variables and model parameters from a previous fit, and only sample-specific latent variables are inferred.

Let us discuss the overall flow of the inference routine in the case mode. First, we instantiate a workspace to keep track of ADVI variational parameters for the continuous latent variables, effective emission potentials, and forward-backward tables and local marginals of all discrete latent variables. We initialize the variational posterior parameters to closely match their corresponding prior distributions. Next, we enter the main stage of inference which consists of interleaved variational updates of the continuous latent variables, and discrete latent variables, each to approximate convergence. Inference over the continuous latent variables is completely delegated to `PyMc3` and is performed using mean-field ADVI algorithm. As mentioned before, our choice of discrete vs. latent variational factorization (Eq. 3) implies replacing the negative binomial log probability with the effective potential given in Eq. (11). We use the Adamax optimizer with a learning rate 0.05 and *β*_1_, *β*_2_ = 0.9, 0.99 to maximize ELBO_C_. We keep track of the downward slope (*m*_*ℒ*_) and the standard deviation (*σ*_*ℒ*_) of the loss function *ℒ* = -ELBO_C_ via exponential moving average smoothing over 500 iterations, and assume convergence once *m*_*ℒ*_ < 0.1 *σ*_*ℒ*_ is satisfied for 10 consecutive iterations. Following approximate convergence in maximizing ELBO_C_, we proceed to the update of the discrete latent variations, i.e. maximizing ELBO_D_. We initialize this step by calculating the effective emission potentials (Eq. 14) via posterior sampling of the continuous variational posterior and evaluating the posterior expectation of the emission potential. We obtain as many posterior samples as needed until a median relative error of < 0.005 is achieved. Next, we calculate the updated marginals of the discrete variables using Algorithm 2 with relaxation coefficient *α* = 0.75 and convergence tolerance *ε* = 0.001. We remark that the forward-backward algorithm involves sequential matrix products along the genomic targets, which we implement efficiently using `Theano`’s sequential `scan` functionality and just-in-time (JIT) compilation.

### Heuristics for avoiding poor local minima

One of the caveats of mean-field variational inference is sensitivity to initialization and propensity to getting stuck in poor local minima. We employ two strategies to mitigate local minimum issues.

#### Marginalized warm-up

It is desirable to marginalize the discrete latent variables *exactly* and perform VI over a model consisting of only continuous variables. Exact marginalization of the discrete variables, however, is intractably difficult. We outlined an approximate VI algorithm based on mean-field decoupling of discrete and latent variables. Such mean-field schemes, however, are particularly sensitive to initialization due to negligence of correlations between the two groups of variables. Intuitively, poor initialization of copynumber posterior leads to co-adaptation of continuous posterior to the wrong copy-number states, which is then perpetuated in the following copy-number posterior updates, and so on. To mitigate this caveat, we have found it useful to marginalize the discrete latent variable *approximately* and perform the first round of *warm-up* ADVI update of the continuous latent variables within such an approximately marginalized model. Explicitly, we neglect the correlations between region-class and copy-number states (i.e. horizontal arrows in **Supplementary Fig. 1c**) momentary. In this simplified model, we can now marginalize both *τ*_*t*_ and *c*_*s,t*_ *exactly*. It is straightforward to show that this recipe amounts to, yet again, simply replacing the negative binomial log probability with the following expression:

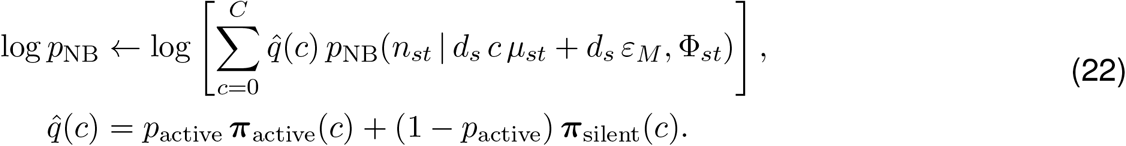

Note the swapped order of summation and log in Eq. (22) and Eq. (11), indicating the non-mean-field nature of the former. After a round of ADVI update of *q*_C_ using such an approximately marginalized model, we proceed to updating *q*_D_. Subsequent interleaved stages of continuous and discrete updates proceed as described before.

#### Entropy regularization (deterministic annealing)

We have found it beneficial to include an entropy regularization term in the loss function. This heuristic is effective for preventing mean-field posteriors from getting stuck in poor local minima due to poor initialization.^1^ In the variational inference framework, entropy regularization can be simply achieved by replacing *H*[*q*] with *T*_*r*_*H*[*q*] in Eq. 1, for *temperature T*_*r*_ > 1. Conversely, one may replace log *p*(**X, Z** | **Θ**) with 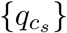. This heuristic is also known as *deterministic annealing*. The original non-regularized ELBO is obtained simply by setting the temperature parameter to 1. In our implementation, we initialize the inference routine with *T*_*r*_ = 2 and linearly anneal the temperature to T_r_ =1 over the course of 500 iterations.

### GATK-gCNV model hyperparameters

GATK-gCNV has two major groups of parameters: those characterizing model-fitting and inference procedures, and those informing the Bayesian priors for the model. The first group includes the optimizer learning rate and maximum number of ADVI iterations, and their default values are set conservatively to ensure convergence of the model-fitting procedure invariably. We determined the default values for key model hyperparameters (*σ*_*m*_, *σ*_*S*_ and *σ*_*T*_) via grid search and identified the hyperparameter combination that resulted in the highest CNV detection F_1_-score over a hold out set from the Simon Simplex Collection samples. **Supplementary Table 2** shows our recommended choice of hyperparameters which were also used throughput this paper.

### PCA-based clustering and batch curation for parallel processing of samples

Exome enrichment kits employ different capture bait designs and as such, exhibit markedly different sequence presentation biases. While GATK-gCNV can learn a general model from sample cohorts encompassing multiple exome enrichment kits, such an application may lead to loss of CNV detection accuracy. For instance, distinguishing highly-polymorphic CNV loci from genomic loci that are subject to high kit-to-kit capture efficiency variability can be challenging. Therefore, it is desirable to avoid grouping together samples that are extracted using different enrichment kits for batch processing. To automate this task, we curated a set of 7,981 target regions chosen from seven commonly-used exome enrichment kits (Agilent_V4, Agilent_V5, Agilent_V6r2, Agilent_V7, NimbleGen SeqCap EZ2, NimbleGen SeqCap EZ3, Illumina TruSeq). Each region was selected to be relatively uniquely represented by one of the seven kits. We collected read-depth over these distinguishing regions across all samples of interest (**Supplementary Table 2**). We normalized the coverage profile by the median sample coverage, performed PCA dimensionality reduction, and clustered samples based using the first three principal components. We ran GATK-gCNV on each sample cluster separately.

### Interval curation using GENCODE v33 exome annotation for CNV detection

In order to harmonize genomic targets across different exome-capture kits that do not all target the same regions, we standardized the set of intervals over which GATK-gCNV queried for CNV events. We took all exons from the GENCODE v33 exome annotation for canonical, protein-coding genes and collapsed them to non-overlapping intervals. Collapsed intervals longer than 1600bp were evenly split into subdivisions less than 800bp, to help increase the number of intervals/data points for GATK-gCNV to identify CNV events. The resulting intervals were symmetrically padded by up to 100 base pairs while ensuring disjointness. The final interval list is provided in **Supplementary Table 3**.

### Filtering intervals for genomic content and sufficient read depth

We excluded intervals with low mappability (<90%), extreme GC-content (<10% or >90%), or prevalent segmental-duplications (>50%), which are known to impede CNV detection accuracy^43–45^. After gathering read-depth information across samples, we further applied a count-based filtering for intervals that are not adequately captured, defined as intervals with a median read-depth < 10 across all samples. We considered the set of intervals passing both filters as well-captured intervals for the purpose of CNV detection across all cohorts.

### GATK-gCNV execution in cohort- and case-modes

GATK-gCNV can be executed in two modes: cohort and case. In cohort-mode, a set of samples are used as input and a model with estimated parameters representing those samples is generated, along with CNV calls. Case-mode takes as input an estimated model generated under cohort-mode and additional samples that are expected to be broadly similar to the ones used to generate the model. Case-mode bypasses the computationally expensive task of model training, and directly infers CNV calls for input samples based on the provided model parameters. We recommend using 100-200 samples for training models in cohort-mode. Aside from using a matching pre-trained model, no sample-size restriction applies to the case-mode execution.

### Postprocessing of raw GATK-gCNV output and calculating CNV quality metrics

Following model-fitting and inference, GATK-gCNV obtains the most-likely sequence of copy number states for each sample using the Viterbi algorithm. The copy number change-points implies a copy number segmentation of the genomic intervals. For each copy number segment, GATK-gCNV generates four quality scores for measuring segmentation accuracy: QA, QS, QSS, QSE. Each quality score is a Phred-scale probability, with larger scores representing higher confidence. These copy number segmentation quality scores were first introduced by the authors of XHMM^10^ and are defined as follows: QA measures the probability that all spanning intervals within the segment are consistent with the copy number assigned to the segment; QS represents the probability that at least one interval is consistent; QSS and QSE represent the probability that the first and last intervals are consistent with being a change-point into and out of the segment copy number state, respectively. In practice, QS provided the highest positive predictive value for progressive filtering and identification of high-quality CNV calls. While the Viterbi algorithm is effective in identifying copy number segments, larger events may be fragmented into several smaller events due to the noisiness of ES data. To remedy this, we extracted non-reference copy number segments from the GATK-gCNV output and considered all segments with quality score QS ≥ 20 for defragmentation (concatenation). We transformed each candidate call into the interval-space and extended each by 50% in width on both ends. We merged segments in the same sample that overlapped after extension and had consistent copy numbers. For defragmented calls, QS was calculated as the maximum QS of the constituent fragments, and QA was calculated as the average. Finally, CNV calls for all samples and clusters were compiled into a single table and single-linkage clustering requiring 80% reciprocal overlap was used to determine when multiple calls were the same site. Site frequencies were assigned to each site as the proportion of samples that had the site in the combined dataset. We also calculated the number of well-captured intervals that each variant spanned, as well as the number of well-captured exons curated from GENCODE v33 annotation described above that each spans.

### Filtering raw GATK-gCNV output for association-level precision and recall

There are two dimensions of filtering, one at a variant-level and one at a sample-level, that we can leverage to produce filtered call sets at desired performance threshold and to meet the desired trade-off between recall and precision. For the variant-level filtering, we found the QS score of a variant to be highly correlated with the confidence of said variant - the higher the QS score, the more likely a variant was found to have orthogonal support in our benchmarking dataset. Our recommended QS filtering thresholds for association studies is as follows:

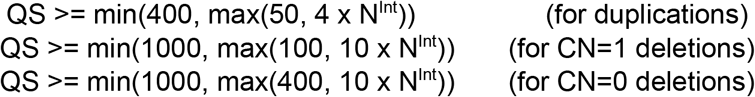

where N^Int^ denotes the number of well-captured intervals that a variant spans. For sample-level filtering, we found the number of raw autosomal GATK-gCNV variants and the number of variants with QS > 20 to be good proxies for how well a sample conforms with the GATK-gCNV model. For association-level filtering, we recommend restricting to samples that have no more than 200 autosomal GATK-gCNV variants and no more than 35 variants with QS >20.

### Benchmarking GATK-gCNV in ASD samples

We had access to 8,439 samples for which matching GS and ES data were available for benchmarking comparisons. Ground-truth CNVs were called from GS using the ensemble machine-learning method GATK-SV^47^. We removed 477 outlier samples that had more than 16 rare (site frequency < 1%) CNVs in the GATK-SV GS ground-truth data, a threshold drawn based on the median + 2 x inter-quartile range of the GS variant distribution, leaving 7,962 samples for comparison. After removing samples that did not pass GATK-gCNV exome QC filters described above (n=927 samples, 11.6%), 7,035 samples remained. Benchmarking was carried out across these 7,035 samples for all rare CNVs (site frequency < 1%). Sensitivity was measured by the proportion of sites called from GS data that have a match in the GATK-gCNV callset. Specifically, for each site, if at least 50% of the samples that have that CNV in the GS data also had a GATK-gCNV call with a consistent direction (deletion or duplication) that overlapped at least 50% of the captured exons, this was considered a success. For CNVs called by GATK-gCNV, their precision was measured by requiring that 50% of the GATK-gCNV samples with that call have a match in the GS calls (ground-truth) with at least 50% interval overlap. Performance was also measured as a function of the number of well-captured exons that a variant spans.

For measurement of the recall of GATK-gCNV ES calls relative to high confidence CMA data, we began with a set of 7,636 samples with matched CMA and ES data. After restricting to large (>50 kilobases & >2 exons), high-confidence (CMA probability < 10^−9^) CNVs from CMA, we found that GATK-gCNV filtered even at our recommended association-level QS thresholds achieved 97% recall of these CMA events, as measured by 50% coverage on the well-captured interval scale in 50% of the same samples.

### Benchmarking XHMM and CONIFER in ASD samples

Out of 7,962 samples that we evaluated for GATK-gCNV, we could access raw sequencing CRAMS for 7,665 samples. We processed these 7,665 samples with XHMM and CONIFER, using the same batches that we created for GATK-gCNV, to minimize the impact of pre-batching on algorithm performance. All CNV tools received as input the list of 330,526 intervals (which corresponded to 190,488 exons from GENCODE v33) that passed GC-content, mappability, and segmental-duplication filtering. Sample- and call-level filtering were conducted according to published best-practices for each method. For XHMM, filtered intervals and failing samples were automatically reported by the method. For CONIFER, filtered intervals were reported automatically by the method, and based on recommended procedure, samples were removed based on CONIFER reported SD value after visualization of the distribution (SD > 0.7).

All XHMM calls across the batches were clustered via single-linkage clustering to determine which calls were the same variant across batches, and the same was done for CONIFER. Frequency of each variant was annotated as the number of carriers divided by the total of passing samples for each method respectively. Using the same definition of recall and precision defined for measuring the performance of GATK-gCNV against ground-truth GS CNV data, we measured the performance of GATK-gCNV, XHMM, and CONIFER against GS CNVs. However, for recall, we measured performance as a function of the number of overlapping, unfiltered exons (190,488), to account for the differential filtering of exons by each of the three methods.

### Benchmarking GATK-gCNV in UKBB samples with CMA data

Owen *et al*.^*32*^ reported population rates of 49 GDs based on approximately 500k UKBB samples using CMA data. We took those 49 GDs and labeled our GATK-gCNV variants as belonging to one of those 49 loci using 50% reciprocal overlap criterion. After harmonizing to the same 49 GDs, we measured the concordance of the CMA derived frequencies in Owen *et al*. to the GATK-gCNV ES derived frequencies, which achieved a correlation of R^2^=0.95, with p-value=1.5×10^−23^, indicating high concordance.

We also used GenomeSTRiP IntensityRankSumAnnotator (IRS^12^) to estimate *in silico* validation rate of copy number variants in the UKBB GATK-gCNV callset using SNP array intensity data. GenomeSTRiP IRS needs two main input files: a VCF file with copy number calls to validate and a matrix of array intensity values. First, we reformatted the GATK-gCNV UKBB output to a VCF file format, containing an *INFO* tag called *SAMPLES* on each site to indicate the set of samples that carry the variant, as required by GenomeSTRiP IRS. Then, we created a matrix of array intensity values for the 177,158 common samples that we could match up between the GATK-gCNV callset and the UKBB SNP array data. This matrix consisted of a tab-delimited text file with four fixed columns (ID, CHROM, START, END) and 177,158 additional columns, one per sample that had array data. Each array-probe location was presented with a single intensity value per sample, the *log R ratio* (LRR), which corresponds to the log_2_-transformed values of the normalized intensity for a given SNP ^46^. LRRs were calculated from UKBB converted VCF files using MoChA (https://github.com/freeseek/mocha/), which calls the gtc2vcf (https://github.com/freeseek/gtc2vcf) plugin. We ran the GenomeSTRiP IRS annotator without using genotypes (*-irsUseGenotypes* parameter set to False) in the Google Cloud. All the scripts used for running IRS, reformatting the VCF file and creating the matrix of array intensity values can be found in https://github.com/talkowski-lab/cnv-validation. GenomeSTRiP IRS outputs an annotated VCF and a report file containing two p-values generated for each site, a negative-shift test for samples with copy number less than the reference ploidy (LOWER p-value) and a positive-shift test for samples with copy number greater than the reference ploidy (UPPER p-value). We assessed CNV sites that have support in the SNP array intensity data for both high- and low-quality calls in the autosomal chromosomes. Since quality metrics are given per site and sample by GATK-gCNV, high quality calls were defined as sites with more than 50% of samples called as high quality (pass sample, frequency and quality filters), and low-quality calls where those with more than 50% samples with a quality score (QS) <= 5. To evaluate validation rates, we kept variants with an allele frequency between 0.01% - 1% and calculated the ratio of the number of sites with a p-value <= 0.1 to the number of sites with a valid p-value for each IRS p-value (UPPER or LOWER) at different minimum number of exons (**Supplementary Fig. 5**).

### GATK-gCNV CNVs in UKBB measured against orthogonal constraint

We curated a set of 16,264 genes based on GENCODE v33 annotation, and which had matching LOEUF scores from gnomAD_v2.1.1 as well as pTS and pHI scores calculated in Collins *et al*. For deletions, we counted the number of rare, high-quality CNVs from the UKBB ES callset that overlapped each of the genes. We then normalized the aggregated count by the number of total coding megabases annotated for that gene in GENCODE V33. For duplications, we carried out a similar calculation, calculating the number of duplications per coding megabase. Lastly, for IEDs, we first identified a set of putative IEDs, defined as duplication events where the two exons on both ends of that gene remain unduplicated, and then counted the number of such events that impacted each gene. Furthermore, due to the difficulty in ascertaining exact breakpoints of events from ES, we only considered 4,634 genes with a relatively large number of exons (>=14, defined as two exons on either end and ten exons in the middle). We normalized the aggregated count again by the number of total coding megabases of the chosen genes. We subsequently compared the normalized count of deletions, duplications, and IEDs against other measures of genic constraint: LOEUF, pTS, and pHI.

### Phenotype association

We extracted 529 traits found to be high-quality by the Pan-UKB team^35^, removing 51 phenotypes that were labeled as continuous but had less than 10 unique values. After restricting to European ancestry, we retained 179,409 samples. We tabulated the overlap of genes by ultra-rare (frequency < 0.1%), high-quality deletions and duplications that ranged in size between 5 and 500 exons and were not categorized as GD CNVs. For each gene count-phenotype pairing, where the number of deletion overlapping that gene numbered at least 5, we executed the SKAT test, adjusting for the top 20 principal components, sex, and age. This resulted in 2,296 tests of gene deletions to phenotype. Similarly, we executed the same SKAT test for duplications against genes with at least 5 duplications, resulting in 4,867 tests. Lastly, we examined all GDs with at least 5 events, and executed the SKAT test with the same covariate adjustments on the 478 traits, resulting in an additional 46 tests. The significance threshold was thus derived as 0.05/478/(2,296+4,867+46)=1.5×10^−8^. We retained association results where the significance threshold was met, and we have at least 5 observations in the exposed group.

---

**Supplementary Fig. 1.**
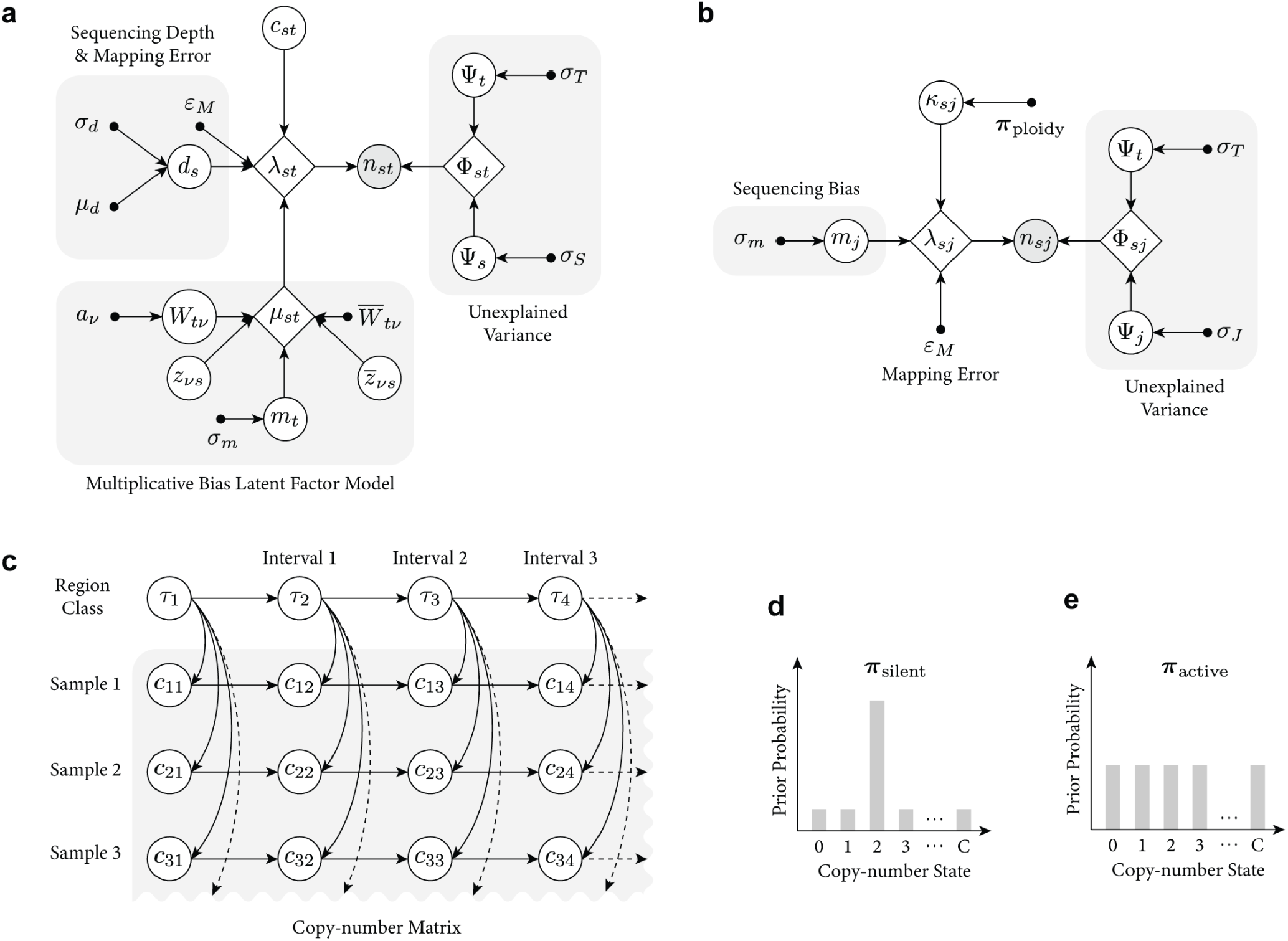
Overview of the likelihood read-depth model and hierarchical Hidden Markov Model (HHMM) that underpins GATK-gCNV. **a**, The likelihood model of read-depth n_st_ at a given interval t and sample s; **b**, The chromosomal baseline copy number 𝒦_sj_ model at given chromosome j and sample s. Observed variables are shown as shaded circles, latent variables as blank circles, hyperparameters as black dots, and deterministic variables as rhombuses. Full parameter definitions appearing in **a** and **b** are given in **Online Methods. c**, The HHMM portion of the GATK-gCNV model consists of primary and secondary chains. The primary chain encodes information on whether the interval in question is a copy number polymorphic site, while the secondary chains encode the actual copy number state of a particular sample at that interval. **d**, The copy number prior distribution that is used to inform the secondary chain for intervals that are predominantly copy-neutral. **e**, The copy number prior distribution that is used to inform the secondary chain for intervals that are copy number polymorphic. **Abbreviations:** HHMM - hierarchical Hidden Markov Model

**Supplementary Fig. 2.**
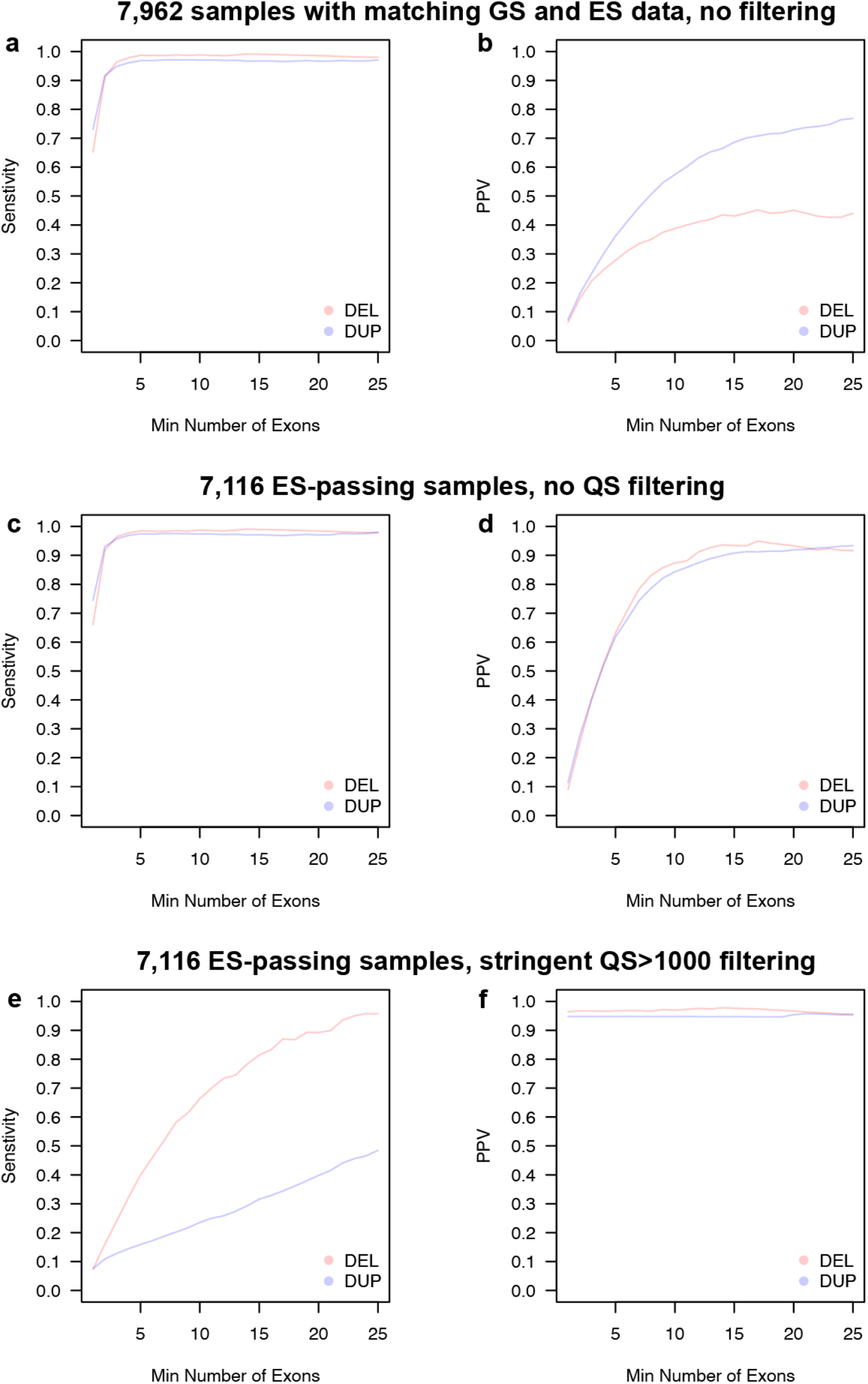
When considering rare CNVs (site frequency < 1%) in 7,962 SSC samples with matching ES and GS data that span more than two exons, raw GATK-gCNV output achieved **a**, 95% recall and **b**, precision of 22%. When restricting to ES-passing samples, we found **c**, 96% recall and **d**, 40% precision at the same size threshold without filtering on QS. Further implementing a stringent QS threshold of >1000, GATK-gCNV produced **c**, precision of 96% for all variants and **d**, sensitivity of 17% at more than two exons. **Abbreviations:** GS - genome sequencing; ES - exome sequencing.

**Supplementary Fig. 3.**
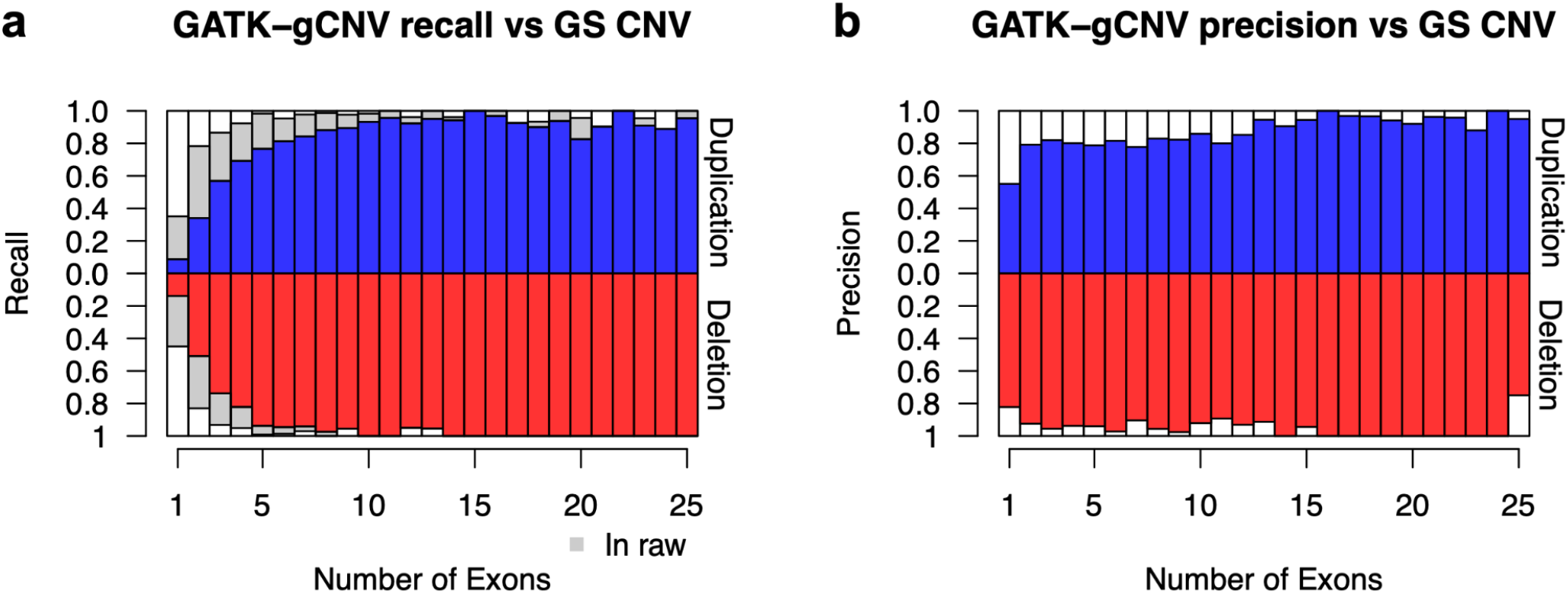
**a**, The recall (and **b**, precision) of rare CNVs in GATK-gCNV ES CNVs compared to GS gold-standard CNVs as a function of the number of exons each variant spans. **Abbreviations:** GS - genome sequencing.

**Supplementary Fig. 4.**
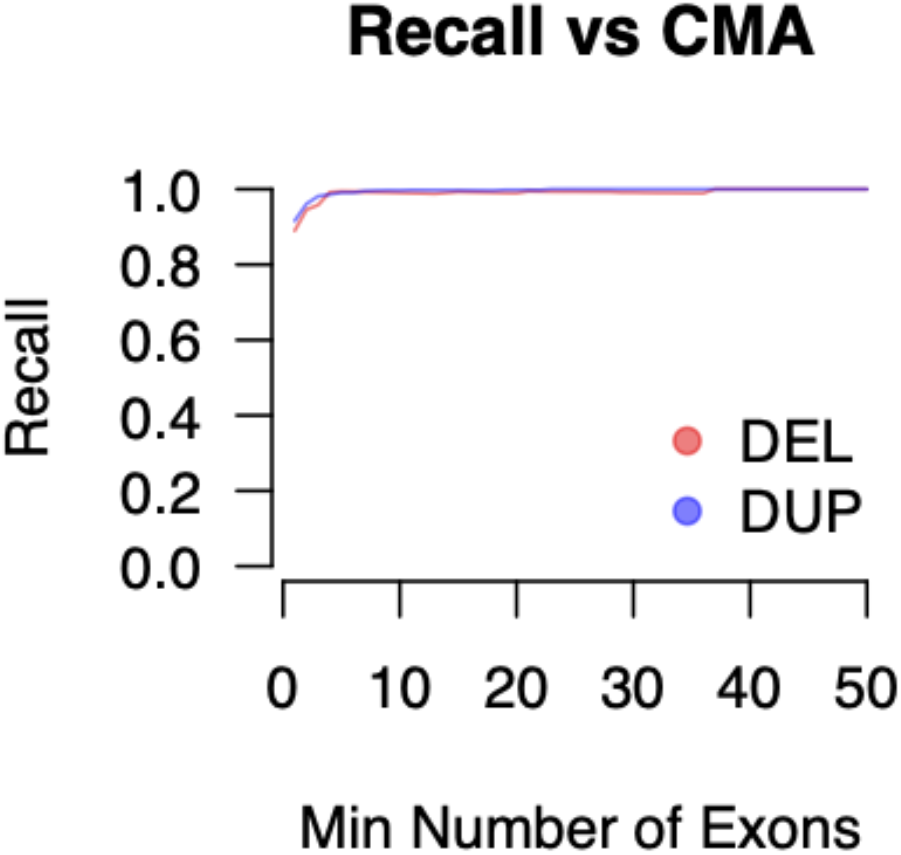
We evaluated the performance of our high-quality GATK-gCNV callset versus rare CNVs (<1% site frequency) identified by CMA in 7,157 SSC samples with matching ES and CMA data. After restricting to large (>50 kilobases & >2 exons), high-confidence (CMA probability < 10^−9^) CNVs from CMA, we found that our high-quality GATK-gCNV callset achieved 97% recall of these CMA events. **Abbreviations:** CMA - chromosomal microarray.

**Supplementary Fig. 5.**
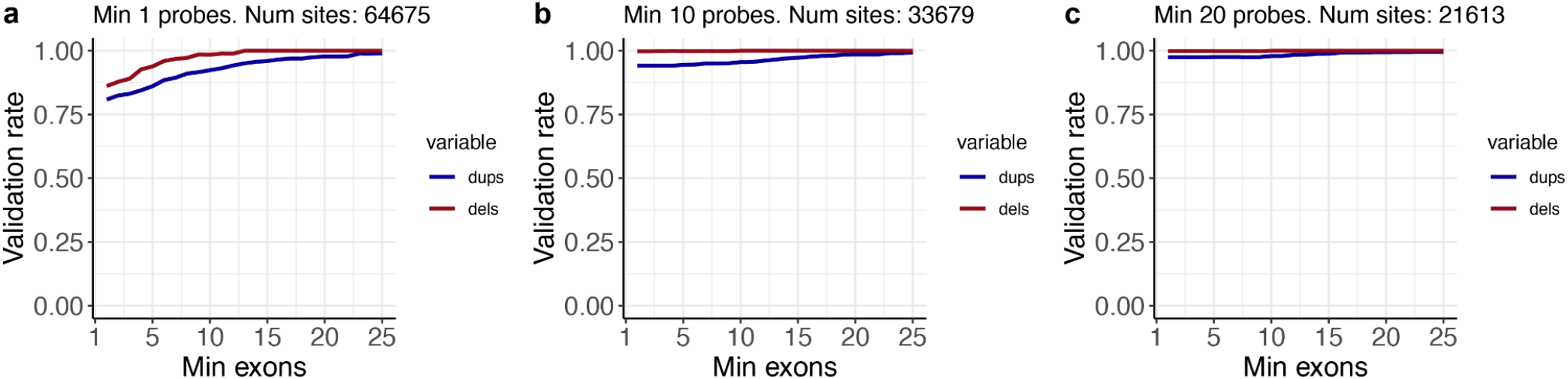
UKBB GATK-gCNV validation rates at different minimum number of exons. Validation rate is calculated as the ratio of the number of sites with a p-value <= 0.1 to the number of sites with a valid p-value for each IRS p-value at **a**, 1 **b**, 10, and **c**, 20 probes, at different minimum number of exons. Only variants with an allele frequency between 0.01% - 1% were evaluated.

**Supplementary Fig. 6.**
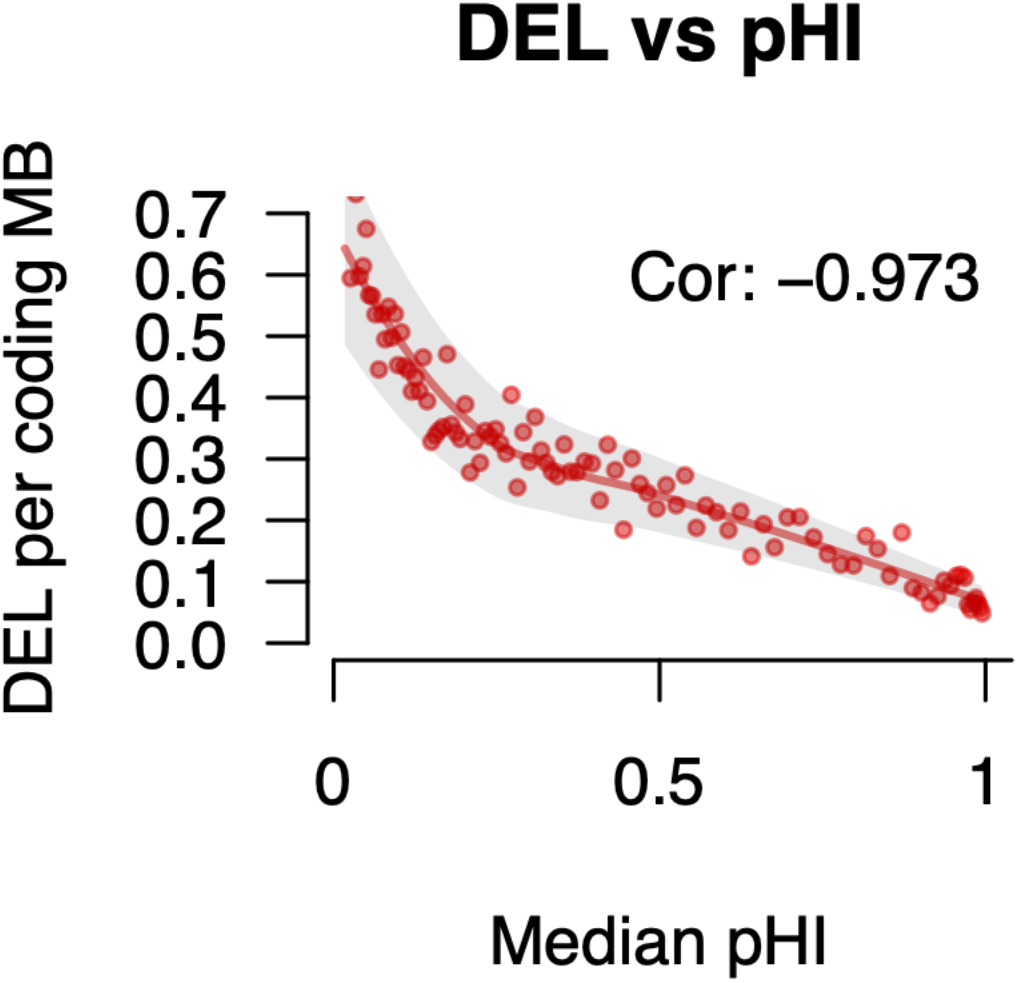
We observed severe depletion of high-quality deletions in our GATK-gCNV callset that overlapped constrained genes as measured by pHI score. **Abbreviations:** DEL - deletion.

**Supplementary Fig. 7.**
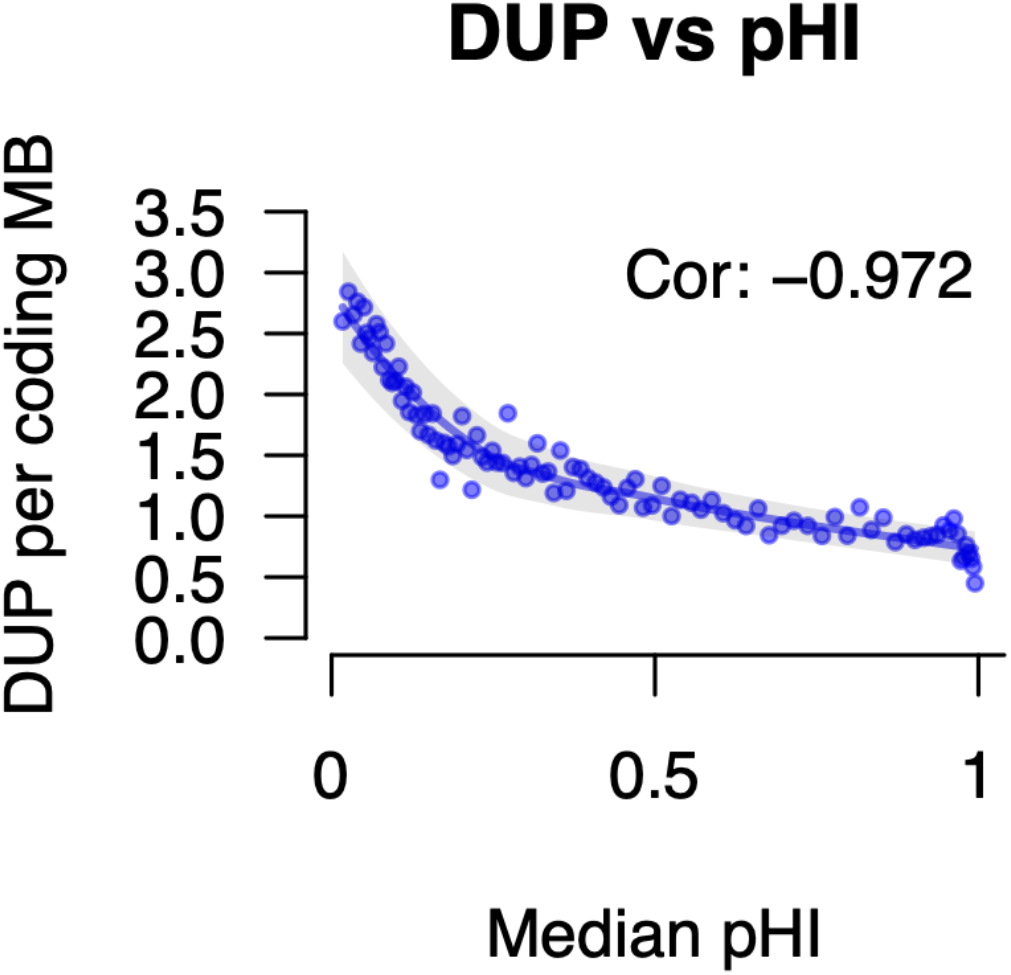
We observed severe depletion of high-quality duplications in our GATK-gCNV callset that overlapped constrained genes as measured by pHI score. **Abbreviations:** DUP - duplication.

