## Supplementary Table 2 for "GATK-gCNV: A Rare Copy Number Variant Discovery Algorithm and Its Application to Exome Sequencing in the UK Biobank"

| GATK-gCNV Tool Argument | Symbol | Type | Default Value |
|---|---|---|---|
| Model hyperparameters | | | |
| `--p-alt` | $p_{\text{alt}}$ | `float` | `5e-04` |
| `--p-active` | $p_{\text{active}}$ | `float` | `0.1` |
| `--cnv-coherence-length` | $d_{\text{CNV}}$ | `float` | `1e+04` |
| `--class-coherence-length` | $d_{\tau}$ | `float` | `1e+04` |
| `--max-copy-number` | $C$ | `int` | `5` |
| `--max-bias-factors` | $D$ | `int` | `6` |
| `--mapping-error-rate` | $\varepsilon_{\text{M}}$ | `float` | `1e-02` |
| `--interval-psi-scale` | $\sigma_{\text{T}}$ | `float` | `1e-02` |
| `--sample-psi-scale` | $\sigma_{\text{S}}$ | `float` | `1e-02` |
| `--log-mean-bias-standard-deviation` | $\sigma_m$ | `float` | `1e+01` |
| `--depth-correction-tau` | $\sigma_d^{-2}$ | `float` | `1e+04` |
| `--num-gc-bins` | $N_{GC}$ | `int` | `20` |
| `--gc-curve-standard-deviation` | N/A | `float` | `1.0` |
| Model-fitting and inference hyperparameters | | | |
| `--learning-rate` | N/A | `float` | `5e-02` |
| `--adamax-beta-1` | $\beta_1$ | `float` | `0.9` |
| `--adamax-beta-2` | $\beta_2$ | `float` | `0.99` |
| `--log-emission-samples-per-round` | N/A | `int` | `50` |
| `--log-emission-sampling-median-rel-error` | N/A | `float` | `5e-03` |
| `--log-emission-sampling-rounds` | N/A | `int` | `10` |
| `--max-advi-iter-first-epoch` | N/A | `int` | `5000` |
| `--max-advi-iter-subsequent-epochs` | N/A | `int` | `100` |
| `--min-training-epochs` | N/A | `int` | `10` |
| `--max-training-epochs` | N/A | `int` | `100` |
| `--initial-temperature` | N/A | `float` | `2.0` |
| `--num-thermal-advi-iters` | N/A | `int` | `2500` |
| `--convergence-snr-averaging-window` | N/A | `int` | `500` |
| `--convergence-snr-trigger-threshold` | N/A | `float` | `0.1` |
| `--convergence-snr-countdown-window` | N/A | `int` | `10` |
| `--max-calling-iters` | N/A | `int` | `10` |
| `--caller-update-convergence-threshold` | $\epsilon$ | `float` | `1e-03` |
| `--caller-internal-admixing-rate` | $\alpha$ | `float` | `0.75` |
| `--caller-external-admixing-rate` | N/A | `float` | `1.0` |
| Miscellaneous hyperparameters and tool arguments | | | |
| `--init-ard-rel-unexplained-variance` | N/A | `float` | `0.1` |
| `--enable-bias-factors` | N/A | `boolean` | `true` |
| `--active-class-padding-hybrid-mode` | N/A | `int` | `50000` |
| `--disable-sampler` | N/A | `boolean` | `false` |
| `--disable-caller` | N/A | `boolean` | `false` |
| `--disable-annealing` | N/A | `boolean` | `false` |

***Supplementary Table 2***. *GATK-gCNV default hyperparameters.*
